# Mass spectrometry imaging-based explainable machine learning reveals the biochemical landscapes of the mouse brain

**DOI:** 10.1101/2025.09.12.675752

**Authors:** Jacob Gildenblat, Jorunn Stamnæs, Jens Pahnke

## Abstract

Recent computational advances in mass spectrometry imaging (MSI) now enable unprecedented insight into organ-wide molecular composition and functional architecture. Here, we present the first high-resolution molecular-computational atlas of specific mouse brain lipids and metabolites, acquired using a NEDC matrix and negative-mode MSI, covering 123 anatomically defined regions and 191 polygonal annotations derived solely from MSI data, without auxiliary imaging. To overcome annotation ambiguity and MSI complexity, we introduced the *Computational Brain Lipid Atlas* (CBLA), a graph-based visual-explainability framework that generates *Virtual Landscape Visualizations* (VLVs) of specific lipid distributions across brain substructures. The CBLA integrates dimensionality reduction and ensembles of supervised models to (i) refine annotations, (ii) elucidate interregional relationships, (iii) interpret model behavior, and (iv) formulate biologically testable hypotheses. The CBLA revealed novel lipid distribution patterns, functional integrations, anatomical connections – the brain’s telephone cables, and region-specific disease signatures – index lipids, including disease networks in the basal ganglia. It further identified *index lipids* that trace extrapyramidal nuclei and their cortical-brainstem connections, highlighting network-level molecular organization. A new algorithm decomposes annotated regions into precise *m/z* features and resolves high-resolution *m/z* values from MSI data, producing a comprehensive high-resolution brain map. It can be applied to any MS measurements, including metabolites, lipids, and peptides. This resource underpins down-stream studies, as exemplified here by characterizing the molecular lipid composition of A***β*** plaques, their spatial arrangement, and their connections with surrounding tissue. For the first time, our data suggest that GM3 accumulation in cortical amyloid plaques may originate from hippocampal structures, consistent with longstanding evidence of disrupted hippocampocortical connectivity; a similar origin may also apply to plaque-associated A***β*** signals in the cortex. More broadly, several selected *m/z* signals showed putative anatomical origins in specific brain subregions.

**Highlights:**

1. Mass spectrometry imaging (MSI) data were used to generate high-resolution, truthful visualizations for brain-region annotation without additional modalities.
2. MSI data were further used to build a computational atlas of annotated brain regions.
3. Pathological structures reveal both their origins and effects on specific brain networks.
4. Anatomical regions and functional networks exhibit distinct lipid/metabolite patterns.
5. Brainstem nuclei and white matter exhibit distinct lipid/metabolite compositions, indicating their involvement in pathological networks.
6. Atlas-based Virtual Landscape Visualizations (VLVs) enable comparison of region-specific differences across mouse models.
7. Several plaque-associated *m/z* signals, including GM3-related species, show putative hippocampocortical anatomical origins.
8. Extrapyramidal nuclei and their cortical-brainstem connections are characterized by shared index lipids, enabling network-level molecular tracing.

## 1 Introduction

Studying brain composition is fundamental to advancing our understanding of its structure, function, development, and disease. Traditional approaches, such as histology, immunohistochemistry, and bulk biochemical analyses, have provided critical insights into neural processes, such as synaptic transmission, energy metabolism, and cell signaling, by probing the distribution of lipids, proteins, and metabolites. However, these methods often lack the spatial resolution or molecular specificity required to fully capture the intricate organization of the brain.

To address these challenges, we used Mass Spectrometry Imaging (MSI) to investigate brain lipid composition. Specifically, we aimed to generate and visualize a highly detailed molecular map of functional brain regions using MSI data alone. To achieve this, we employed recently developed computational methods for dimensionality reduction and MSI visualization [1, 2]. Because MSI data are inherently high-dimensional, both visualization and annotation are challenging. Supervised machine learning (ML) methods require region-level annotations; however, generating such annotations is difficult without clear visual delineation of subregions. This limitation has contributed to the scarcity of prior studies that apply supervised annotation *directly* to MSI data. For example, Huang et al. [3] performed binary annotation (hippocampus versus cortex) on PCA visualizations to compare lipid distributions. Guo et al. [4] relied on automatic segmentation to identify regions, whereas Sun et al. [5] used hematoxylin-eosin (H&E)-stained tissue to generate annotations that were later registered to MSI data. More recently, Xie et al. [6] overlaid MRI-defined annotations onto MSI to create a molecular map of 11 region types. In MSI workflows, immunohistochemical stains (IHC) are often recommended as overlay modalities [6], including MALDI-IHC approaches [7]. In our view, a key limitation lies in how MSI ion images are often presented and interpreted, with insufficient morphology-guided context. Neuropathologists and pathologists can retrieve substantial structural information directly from panels of truthful MSI visualizations [2], without relying on additional modalities.

Moreover, external modalities such as MRI or H&E/IHC are often unavailable, and when used, they can limit annotation granularity to features visible in those modalities. Many brain substructures may not be distinguishable with pathology stains such as H&E, which are blind to lipids, nor with MRI alone. While Schede et al. [8] recently demonstrated the feasibility of MSI for profiling approximately 100 hypothetical lipids through lipid-database mapping of *m/z* values in *Danio rerio* fish embryos, their analysis made only limited use of spatial information. In contrast, our approach enables high-fidelity mapping of lipid distributions across a broad range of anatomically defined brain regions. Similar to Schede et al., we generated mappings of *m/z* values across regions; however, by focusing specifically on the imaging dimension of MSI and truthful visualizations in a multi-visualization panel [1, 2], we were able to resolve a much larger number of regions and, crucially, provide new tools for studying their relationships and compositional organization.

Here, we show that MSI contains rich, underused information that can support fine-grained expert annotation and supervised ML. We present MSI-ATLAS, a customizable pipeline for integrated annotation and analysis of high-dimensional MSI data (e.g., metabolites, lipids, and peptides) without relying on external imaging modalities. To support structural annotation, we developed MSI-VISUAL, an actively developed framework that combines multiple MSI visualization methods, including established dimensionality reduction approaches, to generate detailed, information-rich images [2]. This multi-visualization strategy reveals subtle anatomical structures that are often missed by conventional segmentation and therefore supports human-guided polygonal annotation of regions and subregions. In this study, we combine MSI-VISUAL-based annotation with large-scale mapping of *m/z* values associated with each brain region using two complementary computational approaches. To resolve *m/z* features at high precision, we additionally employed a custom Gaussian Mixture Model-based approach.

A central challenge in this work is the visual exploration and interpretation of lipidomic relationships between brain regions. Because brain organization is complex and MSI data are high-dimensional, regional annotations can partially overlap and categories may occasionally be ambiguous or mislabeled. To evaluate interregional molecular relationships, we developed visualization tools that interrogate the distribution of features (*m/z* values) within and between annotated regions and categories. These tools also support diagnosis and interpretation of machine learning models trained on the data. For interactive exploration of the lipid landscape, we developed the *Computational Brain Lipid Atlas* (CBLA), an interpretable graph in which each annotated region or category is represented as a node, with edges reflecting biological or annotation-based molecular similarity. The CBLA can be overlaid with *Virtual Landscape Visualizations* (VLV), which represent the response of annotated regions to specific *m/z* values. Together, these tools allow users to query *m/z* values and examine their distribution across gross anatomical regions and finer subregions, enabling the generation of biologically testable hypotheses from MSI data.

As an application example, we use MSI-ATLAS to characterize lipid complexity in the brain by generating 191 polygon annotations mapped to 123 brain region types according to the Allen mouse brain atlas [9]. To our knowledge, this is the first comprehensive molecular map of brain substructures using human-labeled MSI data at this level of detail. We find that anatomically and functionally related regions share characteristic lipid composition profiles. Brain connectivity is reflected in molecular structure: functionally connected regions, such as extrapyramidal brainstem nuclei, exhibit shared lipid signatures, resembling a colored wiring diagram. Finally, we demonstrate the utility of this pipeline by analyzing A*β* plaques in a mouse model of Alzheimer’s disease and characterizing their molecular relationships with surrounding brain regions, revealing distinct plaque-associated lipid signatures linked to affected cortical regions.

## 2 Materials and Methods

### 2.1 MSI Data Generation and Extraction

#### The MSI mouse brain dataset

We generated mouse-brain MSI data from frozen hemispheres of four 100-day-old C57Bl/6J animals. Two mice expressed combined human APP and PS1 mutant transgenes (APPtg) for A*β* plaque formation [10–12], and two mice were ABCA7 knockouts [13]. Frozen hemispheres were cut into 10*µm*-thick coronal sections, before and after bregma [14], using a cryomicrotome (CM1950, Leica, Nussloch, Germany) and placed on a single Intellislide (Bruker Daltonics, Bremen, Germany). Sections were dried in a vacuum chamber and sprayed with N-(1-Naphthyl) ethylenediamine dihydrochloride (NEDC) matrix (7 mg/ml in 70% methanol) at 30^◦^C using an HTX3+ TM Sprayer (HTX Technologies, Carrboro, NC, USA), adapted from Andersen et al. [15]. Matrix-assisted laser desorption ionization mass spectrometry imaging (MALDI-MSI) was performed in negative mode using a timsTOF fleX™ mass spectrometer (Bruker Daltonics, Bremen, Germany). We performed mass calibration using red phosphorus. Ion mobility was calibrated using ESI-L Low Concentration Tuning Mix (Agilent Technologies, Santa Clara, CA, USA) in ESI mode; however, ion-mobility information was not used for feature extraction or downstream analyses in this study. For MSI analysis, we used a mass range of *m/z* 300–1350 Da. The laser operated at 10 kHz with a burst of 200 shots per pixel at a spatial resolution of 20 *×* 20*µm*^2^. We used the machine standard of 20 *×* 20*µm*^2^ (4 *×* 4 pixels of 5 *×* 5*µm*^2^) to generate sufficient signal (AU), including low-abundance species. A 5 *×* 5*µm*^2^ setting provided higher spatial resolution at the cost of reduced low-abundance signal. Four mouse hemispheres were assessed in a single overnight run on one slide.

#### Data extraction from MSI files

We used the MSI-VISUAL toolbox [2] to extract data in the measured *m/z* range of 300–1350 Da with fixed-bin resolutions of 5 and 20 bins per *m/z* (0.2 and 0.05 Da, respectively). For each image pixel, detected *m/z* values were quantized to the nearest bin, and their intensities were summed into that bin. We used the 5-bin extracted files for visualization, whereas 20-bin extracted files were used for initial statistical comparisons before high-precision refinement with Gaussian Mixture Models (GMM). High-resolution *m/z* values were determined from the *raw* MSI files (see next paragraph). For each extracted 5-bin image file (*n* = 4), we created a multi-visualization panel using MSI-VISUAL [2]. We generated multiple visualizations for each hemisphere using Saliency Optimization (SALO), TOP3, and Percentile Ratio (PR3D), together with dimensionality-reduction methods such as UMAP [16], PCA [17], and non-negative matrix factorization [18]. For subsequent brain-region annotation, we generated a composite overlay of SALO, TOP3, and PR3D visualizations to reveal finer structural details across regions [2]. We also used segmentation methods (e.g., K-means [19]) with targeted visualizations within segments. All visualizations used total ion current (TIC) normalization [20].

#### Resolving high-precision *m/z* values

Brain-region *m/z* values were initially determined at 0.05 Da resolution from a fixed 20-bin representation of the MS images. The goal was to obtain higher-precision *m/z* measurements. We used a four-stage workflow:

1. We re-extracted the original raw MSI spectra, focusing on narrowed *m/z* candidates and recording intensities within a range of up to 0.01 *m/z* around each value. At this stage, we did not generate images; instead, we recorded the *m/z* distributions.
2. For each *m/z* value, we fitted a Gaussian Mixture Model (GMM) [21] using scikit-learn [22] with three components. Minor components representing less than 5% of the data were discarded. Component centers represented *m/z* distributions.
3. We then re-extracted MSI images using these components and summed *m/z* values within a 0.01 *m/z* range around them.
4. Finally, we recomputed *m/z* statistics to verify the high-precision *m/z* values.

### 2.2 Annotation

#### MSI-based annotation of brain regions and subregions

Polygon annotations were performed with the *VGG Image Annotator* [23] by a neuropathology specialist (J.P.), following the Allen mouse brain atlas (AMBA) [9]. Each annotated region was carefully compared across visualizations in the multi-visualization panel (Fig. 1). Four iterative annotation rounds were performed, during which polygon annotations were refined based on assessment using the generated Computational Brain Lipid Atlas (CBLA). Additional subannotations were added to AMBA regions when detectable in the visualizations and were labeled with ‘sub’ followed by an integer. Each annotation was categorized according to AMBA nomenclature into main anatomical categories (gross brain regions), such as cortex (CTX), cerebral nuclei (CNU), brainstem (BST), white matter tracts (WM), choroid plexus (PLX), pathological changes (e.g., A*β* plaque category, PLQ), and background (BG). These main categories were used to color-code resulting data (e.g., nodes) in CBLA diagrams.

**Fig. 1.**
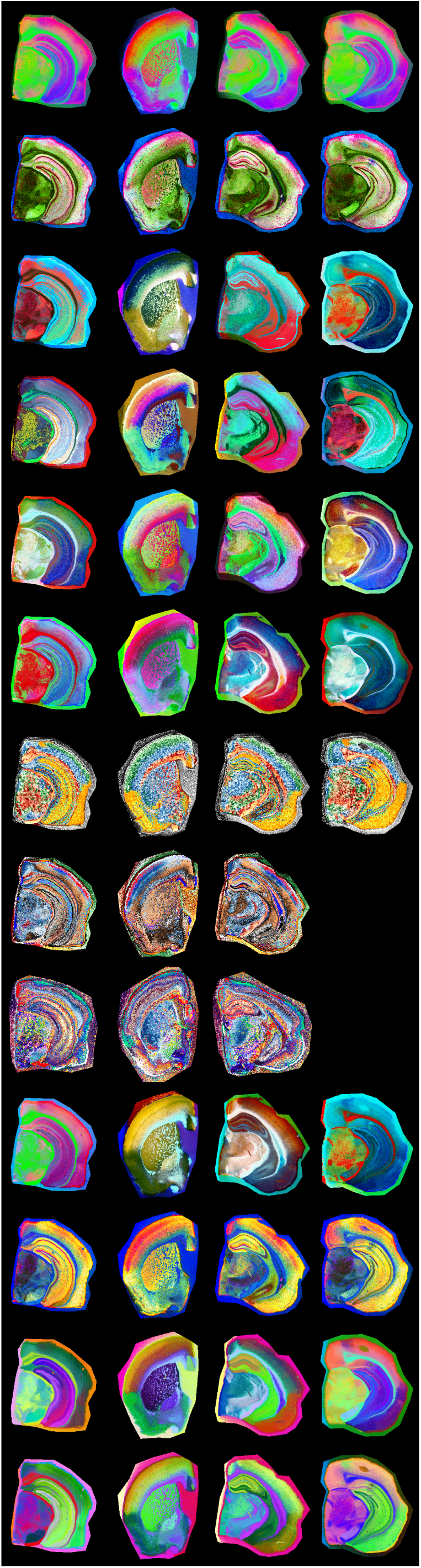
Overview of a multi-visualization panel for brain-region annotation. We used custom-generated panels from four brains at different bregma locations to enable high-resolution annotation of brain regions according to the Allen mouse brain atlas. Each row represents one visualization method generated using MSI-VISUAL [2].

#### Resolving annotation problems using the CBLA

In each annotation iteration, we searched for nodes that were very close to each other or overlapping and connected by strong edges. These nodes were further investigated for their *m/z* profiles to determine unique *m/z* values that could verify both edge representation and node locations in the CBLA.

### 2.3 Modeling

#### Machine learning-based classification of pixel categories

For feature-based pixel classification, we employed a simplified Bayesian linear classification model, implemented with Pyro [24] using logistic regression. Each feature weight was assigned a *Gaussian prior* to encode uncertainty in its contribution to predictive signal. During inference, we approximated the posterior distribution over weights using variational inference. To generate predictions, we sampled weights from the learned posterior distribution and averaged resulting predictions, enabling a probabilistic interpretation of model confidence. Because the model is linear, model weights can be used for interpretability and for identifying significant *m/z* values per region, as described below.

#### Ensemble-based evaluation of category confusion

To quantify worst-case ambiguity between annotated categories, we evaluated category confusion using an ensemble-based strategy. We trained an ensemble of 10 independently initialized models with the Bayesian architecture described above. For each trained model, we estimated the soft confusion matrix by averaging predicted probabilities over 100 stochastic forward passes (sampling from the variational posterior). Each entry in the confusion matrix (*i, j*) represents the average probability that a sample from true category *i* is assigned to category *j*. To assess worst-case confusion across the ensemble, we computed the maximum confusion per pair of categories:

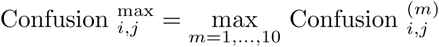

This approach highlights the strongest disagreement between categories observed across model variations and posterior samples, serving as a conservative estimate of label ambiguity and potential annotation or biological overlap.

#### Mapping *m/z* values for brain regions

To identify *m/z* values associated with specific categories (brain regions or subregions), we employed two approaches with distinct behavior: i) a model-based approach that trains an ensemble of models and measures feature importance for *m/z* values, and ii) a model-free approach comparing pairs of brain-region categories to mine *m/z* values that are significant in those categories, denoted *mPAUC*.

Both approaches have different selection behavior. mPAUC tends to select *m/z* values less associated with other categories, whereas model-based selection can also select *m/z* values shared with other brain-region categories, as long as they remain predictive. Supplementary Fig. S1 shows effects of these selection methods for the plaque (PLQ) category. We measured the log_2_ fold-change for PLQ annotations relative to remaining non-background annotations. Note that *m/z* values associated with PLQ may have negative log_2_ fold-change, for example because they are highly abundant across various cortex categories. These *m/z* values may still be associated with plaques and should therefore be selected. The intersection of both selection methods yields *m/z* values that are highly selective for the PLQ category.

#### Model-based selection of *m/z* markers for each brain region

To identify category-specific *m/z* features and interpret linear-model weights, we developed a custom feature-importance scheme inspired by concepts from *boruta* [25] and *STABL* [26]. *Boruta* is a wrapper-based feature-selection method designed for tree-based models, which compares real-feature importance to permuted noise features using random forests. *STABL*, by contrast, is based on sparse linear models and stability selection, combining resampling with noise-informed thresholds to control false discoveries. Although both methods use permuted features and repeated subsampling, they are structurally distinct. Given the computational demands of *STABL* and the tree-based nature of *boruta*, we implemented a lightweight alternative tailored to our linear setting. Specifically, we trained 200 logistic regression models, each on a random 50% subsample of the data. To calibrate feature selection, we introduced *N* = 50 noise features by randomly selecting and permuting existing *m/z* values. For each model and category, we computed the maximum absolute weight assigned to any noise feature and selected real *m/z* features whose positive weights exceeded this threshold. Finally, following the stability-selection principle, we retained only features selected in at least 80% of the models.

#### *m/z*-Pairwise-AUC (mPAUC) – ranking *m/z* values per region by comparing region pairs

This approach assigns a category score for every *m/z* value based on AUC ranking. For each *m/z* value, we scored each category by comparing its separation from other categories and summing AUC values with low p-values. This score indicates that the *m/z* value distinguishes that category from others. We then selected the top 10 categories with the highest score for each *m/z* value. During this comparison, irrelevant categories may also participate and skew the result. To promote competition between relevant categories, we repeated this step using only the top 20 categories for each *m/z* value. In other words, we identified categories scoring highly for each *m/z* value and compared them again without irrelevant categories from the first round. For each *m/z* value *m* and category *i*, we computed a category score by accumulating AUC scores between that category and other categories with p-value below 0.05:

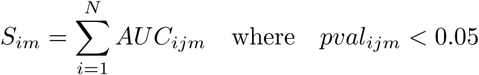

For each *m/z* value *m*, we retrieved categories with the highest *k* = 20 scores and recomputed *S_im_* using this set.

#### Putative annotation

Putative metabolite and lipid annotations were assigned by accurate-mass matching of detected *m/z* values acquired in negative ion mode, primarily assuming [M *−* H]^−^ adducts. Candidate matches were obtained from public databases (e.g., HMDB [27] and LIPID MAPS [28]) using a 0.01 Da mass tolerance, then filtered by lipid-class plausibility, ionization behavior, and known brain biology. Where multiple candidates were possible, assignments were reported at lipid-class or sum-composition level. All annotations (Supplementary Tab. S3) are putative and require orthogonal validation (e.g., MS/MS) for confirmation.

### 2.4 Construction of the Computational Brain Lipid Atlas

The Computational Brain Lipid Atlas (CBLA) is a graph-based framework that enhances interpretability of MSI data, annotations, and trained models (Fig. 3). It provides a unified visual representation supporting exploration of biological structure, annotation reliability, and model behavior.

#### Defining molecular relationships between brain regions

To understand molecular relationships among annotated brain regions, we first computed average *m/z* intensity profiles for each region. We embedded these region-level molecular profiles into two dimensions using UMAP [16]. Each region was visualized as a node positioned according to its 2D embedding. We weighted edges between nodes using maximum confusion values from an ensemble of 10 Bayesian classifiers, with each model’s normalized confusion matrix averaged over 100 posterior samples by taking mean + 2 *· std* of confusion. Maximum pairwise confusion between models was used to define edge weights between nodes (brain regions). Edges between nodes within the same anatomical category (e.g., cortex layers or white matter tracts) were colored red, suggesting likely sources of annotation ambiguity or shared signal, such as biological or functional similarity. Edges between different anatomical categories were colored bright green, indicating potential biological links or molecular similarities that warrant hypothesis generation.

#### Explaining and refining the annotations

CBLA also provides a lens for evaluating annotation quality. Edges with high confusion between regions expected to be distinct (e.g., functionally unrelated areas) suggest potential annotation errors, class imbalance, or poorly defined categories. Conversely, high similarity between nominally different but biologically similar subregions may suggest duplicated or unnecessary labels. This allows researchers to iteratively refine annotations and resolve inconsistencies in dataset structure.

#### Explaining the model

To visualize the model’s internal representation of brain regions, we constructed a parallel version of CBLA using learned prediction weights for each category instead of average *m/z* profiles. These region-level weight vectors represent molecular features the model relies on for classification. Dimensionality reduction and edge construction followed the same process as in the data-based atlas (Supplementary Fig. S2). By aligning data structure, annotation signals, and model decisions, CBLA enables transparent inspection and quality assessment of the MSI-based brain map, supporting both error diagnosis and biological discovery.

### 2.5 Visualizations

#### Colorizing the atlas - Virtual Landscape Visualizations (VLV)

For a selected *m/z* value, we obtain an intensity value for every category node. For data explanation, this value represents average category intensity for that *m/z*. For model explanation, it represents the model category weight for that category. For every pixel, we compute Euclidean distances in coordinate space *d_i_* from all graph nodes (i.e., node 2D coordinates after dimensionality reduction), and compute a weighted average of node representations with a softmax [29]:

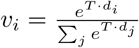

The relationship of the selected *m/z* value with the CBLA network is visualized as a topographic overlay color scheme encoding the 0–1 normalized total ion current of that *m/z*.

#### Visualizing mouse model and region-specific differences with the CBLA

We grouped mice into two cohorts (APPtg+ vs. APPtg– (wild-type), or A7+ (wild-type) vs. A7– (ABCA7 knockout)) and computed average values within each group for every annotated brain category. However, not all brain categories are annotated in every scan, which can prevent computation of group averages for some categories. One way to address this would be to require annotations for all brain categories in each scan. Instead, we adopted a different approach: predicting brain categories using machine-learning-based segmentation. Specifically, we used an ensemble of 100 linear models and assigned categories based on majority vote across the ensemble.

#### Virtual Pathology Stain (VPS)

To simulate a commonly used immunohistochemical (IHC) stain, grayscale ion-intensity images were mapped to a brown color scale. Total ion current (TIC) intensities of fully resolved *m/z* values were normalized to 0–1 for each *m/z* across all pixels, and linear interpolation was performed for each pixel between white (RGB: 255, 255, 255) and dark brown (RGB: 139, 69, 19). This mapping ensures that low-intensity pixels remain light, while high-intensity regions are rendered in progressively deeper brown shades, visually resembling 3,3’-diaminobenzidine (DAB) staining used in histopathology (e.g., Fig. 5c).

## 3 Results

### 3.1 MSI-ATLAS Enables High-Detail Exploration and Mapping

A schematic depiction of the MSI-ATLAS workflow is shown in Fig. 2. MSI-ATLAS was established to generate functional and anatomical representations of organ-specific multi-omics networks. While demonstrated here using brain metabolomics/lipidomics MSI data, the workflow is **generalizable to other spatial omics modalities**, including proteomics.

**Fig. 2.**
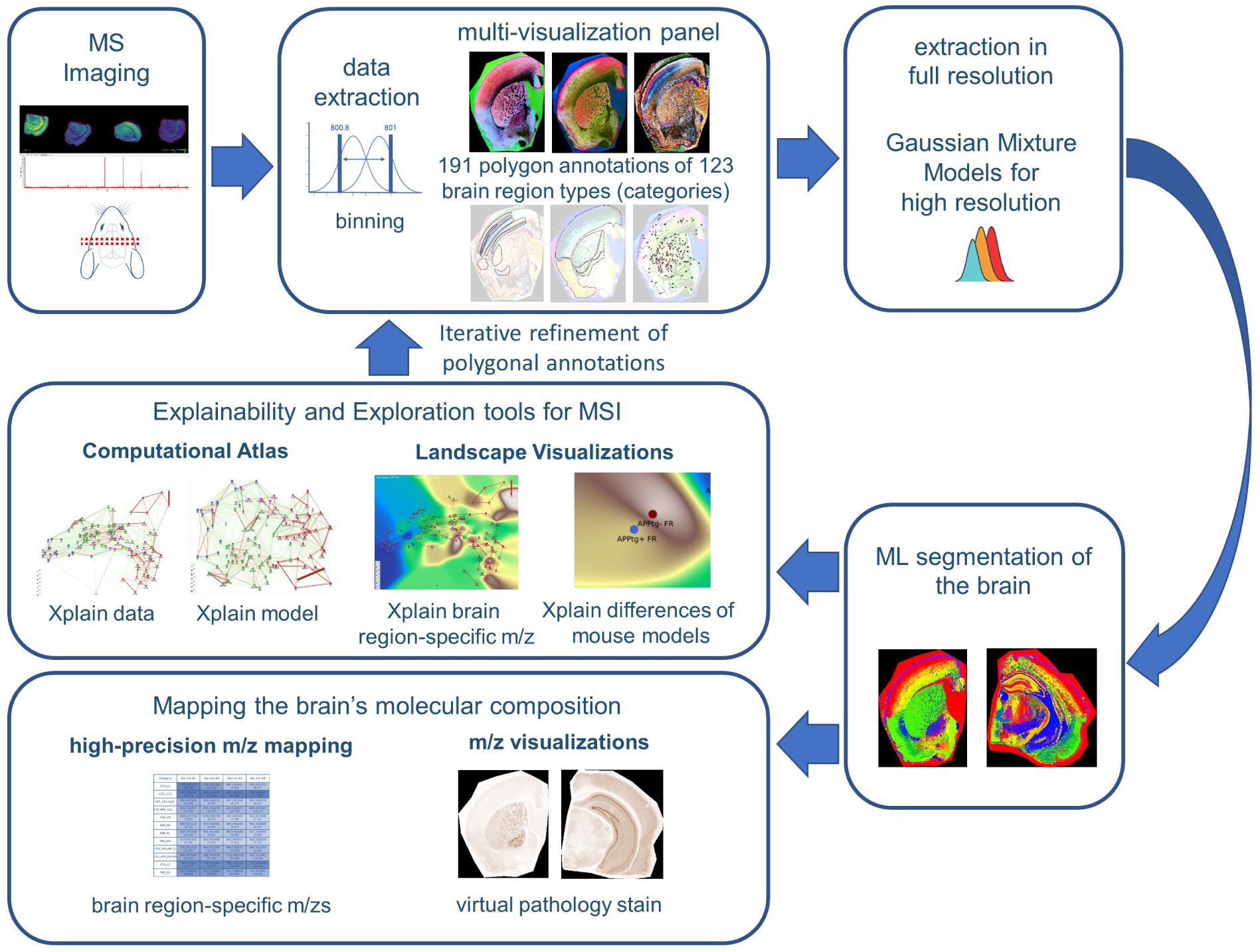
Overview of the MSI-ATLAS workflow. The *MSI-ATLAS* workflow involves comprehensive analysis of mouse brain tissue using mass spectrometry imaging (MSI). The resulting high-dimensional datasets were converted to fixed-bin representations and visualized through advanced visualization methods provided by *MSI-VISUAL* [2]. These reliable visualizations enabled 191 highly detailed polygon annotations based on Allen mouse brain atlas nomenclature [9] and identified newly detected subregions. The resulting dataset comprised 123 brain region types (categories), along with an additional background category, all mapped against binned *m/z* values. Following this mapping, full-resolution extraction from raw mass spectrometric data was performed, and Gaussian mixture models were employed to derive high-resolution *m/z* values. Using the full-resolution data, we created the Computational Brain Lipidomics Atlas (CBLA), where node positions are determined by UMAP dimensionality reduction of average category spectra (brain region types) into two dimensions. This computational atlas was subsequently overlaid with a *Virtual Landscape Visualization* (VLV) to map distributions of selected *m/z* features across various brain regions and to compare mouse models. We used ensembles of Bayesian machine learning models and posterior sampling to compute between-category confusion values that define graph edges. We also used the models to map stable *m/z* values associated with each brain region type. The mapping was visualized for verification using a *Virtual Pathology Stain* (VPS) technique designed to mimic traditional immunostained slides. The CBLA and VLVs were used iteratively to refine annotations.

To establish and evaluate the MSI-ATLAS workflow, we generated and analyzed an MSI lipidomics dataset of four full brain hemispheres from control and APP-transgenic mice representing aged, healthy, and diseased brains, respectively. The dataset comprised 20,020 *m/z* values across four hemispheres at a spatial resolution of 20 *×* 20 *µ*m^2^ per pixel.

We enabled an expert neuropathologist (J.P.) to create region and subregion annotations directly on MSI data by leveraging a multi-visualization panel of complementary views (Fig. 1), each optimized to highlight different molecular patterns [2, 30]. Using these polygonal annotations, we mapped molecular composition for each annotation and created a database of *m/z* values for 123 brain region types. We employed a novel method based on ranking category-specific *m/z* values, *m/z*-Pairwise-AUC (mPAUC), and a machine-learning (ML) model-based approach using feature selection of stable features to associate *m/z* values with brain regions. The resulting region-specific *m/z* associations form a molecular atlas that may serve as a resource for future studies.

To effectively explore and interpret the lipid landscape of the brain, we developed an explainability and visualization tool: the *Computational Brain Lipid Atlas* (CBLA). With 191 annotations for 123 brain region types, we constructed high-dimensional graph representations of each brain region and embedded them as **nodes**. In this graph, **edges** reflect molecular similarity, whether biological or annotation-induced, based on how well ensembles of ML models distinguish brain regions. Background colors in the *Virtual Landscape Visualization* (VLV) represent category responses to specific *m/z* values. CBLA allows users to query *m/z* values and interactively examine their distribution across gross anatomical regions and small subregions. Using CBLA, we observed that anatomically and functionally related regions share characteristic lipid-composition profiles. Brain connectivity appears mirrored in the molecular structure: functionally connected brain regions exhibit shared lipid signatures, akin to a colored wiring diagram. CBLA also supports iterative annotation refinement by identifying regions that may benefit from subdivision into smaller subregions. This feedback loop helped us improve the quality and granularity of our initial annotations and detect previously unknown subnuclei, such as those in the substantia nigra and globus pallidus.

To address the practical challenge of fixed-bin *m/z* representations used in MSI visualizations, we developed a workflow to recover high-precision *m/z* values. Fixed binning can obscure distinct molecular species, especially at high resolution. We modeled distributions of significant *m/z* values with Gaussian mixture models [21], allowing us to identify and refine high-precision molecular features. Finally, the workflow includes tools for spatial *m/z* visualization using the Virtual Pathology Stain (VPS), which mimics commonly used DAB staining for immunohistochemistry, and a novel approach to extract high-accuracy *m/z* values from annotated regions of interest (Fig. 2, Methods and Supplementary Data).

We demonstrate that CBLA provides several biological insights. We used CBLA to study spatial lipid-distribution patterns in A*β* plaques and their relationships with other brain regions, and to visualize region-specific differences across mouse models. We also identified distinct molecular signatures in ABCA7 knockout mice [13].

### 3.2 Construction of the Computational Brain Lipid Atlas

The *Computational Brain Lipid Atlas* (CBLA) is a visual explainability tool built for iterative refinement and exploration (Fig. 3). CBLA was generated after each annotation round, guiding a total of three refinement iterations. As described in detail in the Methods section, the CBLA aggregates MSI data by brain region, applies dimensionality reduction to reveal global organization, and constructs a region-to-region graph based on confusion scores derived from machine-learning classification models. Edge strengths in the graph reflect model uncertainty or misclassification rates, offering insight into potentially overlapping biology or ill-defined regions. Larger nodes represent higher confusion with other categories.

**Fig. 3.**
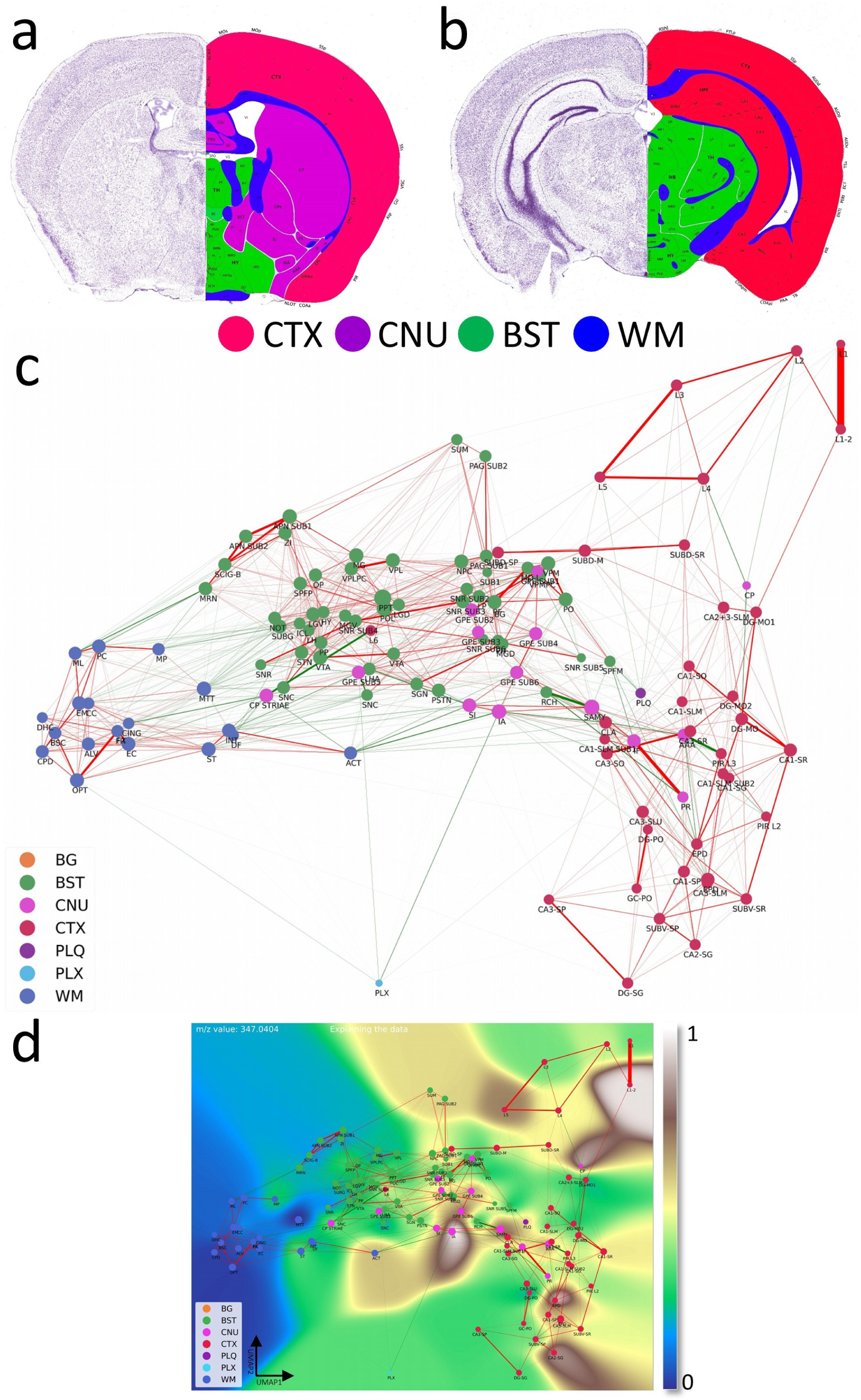
Computational Brain Lipid Atlas (CBLA) and Virtual Landscape Visualizations (VLV): an enhanced analytical framework for molecular profiling. **a,b**, Coronal sections at two fronto-occipital levels, colorized by gross anatomical regions: CTX (red), CNU (violet), BST (green), and WM (blue). This color scheme was used to color-code nodes in the CBLA graph. The left side of each slide shows a Nissl stain of neuronal distribution. The images were adapted from the Allen mouse brain atlas [9]. **c**, The CBLA provides visualization and analysis of brain lipid species (dataset PXD056609), using Virtual Landscape Visualizations (VLVs) to elucidate underlying data structure and ML-model behavior. Graph nodes were constructed by averaging *m/z* spectra across distinct brain-region categories from all measured mice, followed by UMAP dimensionality reduction [16]. Spatial proximity between nodes reflects molecular similarity, facilitating comparative analysis. Node size is proportional to the number of connected edges (confusion) with other categories. Graph edges denote relationships between identified categories derived from classification by multiple ML models and informed by confusion-matrix errors. Green edges indicate connections across different gross anatomical regions, suggesting potential biological associations. In contrast, red edges linking categories within the same gross anatomical region (e.g., cortex layers CTX L2–L5) may indicate annotation ambiguity or intrinsic biological similarity. **d**, The VLV shows the dataset response to the specific *m/z* value 347.04044. Here, nodes represent average spectral values for each category, effectively capturing core data characteristics (dataset PXD056609). Category response to this *m/z* is displayed with the Matplotlib ‘terrain’ colormap [31], enabling interpretation of molecular distributions across brain regions. The 0–1 normalized expression of *m/z* 347.04044 is localized to distinct regions, including cerebral cortex (CTX; red regions corresponding to L1*/*2, CA2-sg, and EPD) and cerebral nuclei (CNU; pink regions corresponding to caudate-putamen [CP], external globus pallidus [GPE], and intercalated amygdaloid [IA] nuclei). Legend: BG, background; BST, brainstem; CNU, cerebral nuclei; CTX, cerebral cortex; PLQ, A*β* plaques; PLX, choroid plexus; WM, white matter.

By introducing *Virtual Landscape Visualizations* (VLVs) of both the data (Fig. 3d) and the model (Supplementary Fig. S2), we enable spatial interpretation of responses to individual or grouped *m/z* values and evaluation of model certainty. The virtual landscape is color-coded by predicted *m/z* response intensity across categories and interpolated to form a smooth, easy-to-interpret map.

Supplementary Fig. S2 provides the model-explanation view used to interpret category weights in the CBLA.

Even before focusing on individual lipids, the CBLA already reflects known brain organization in an intuitive way: nodes from the same gross regions tend to cluster together, whereas anatomically distinct systems are separated in the graph. At the same time, edges between nearby subregions within a gross region (for example, neighboring cortical layers) and select cross-region links are biologically plausible, consistent with shared molecular composition in related structures and selective connectivity between brain systems.

### 3.3 Brain Regions Exhibit Unique Lipid Signatures

First, we used CBLA to understand global organization of brain lipids within three gross anatomical regions: cortex (CTX), white matter (WM), and brainstem (BST). Fig. 4 shows the color-coded organization of 123 mapped brain categories and background as nodes in the CBLA. Nodes from each gross anatomical region cluster together at different graph locations, which agrees with their spatial and functional separation in the brain and resulting *m/z* signatures. Thus, CBLA accurately condenses and visualizes functionally and anatomically distinct regions solely based on MSI data. Nodes that do not cluster with their gross anatomical region represent interesting categories with variant anatomical or functional behavior, e.g., cortex layer 6 (CTX L6), which is located within the central BST cluster, highlighting its molecular nature as a major cortex layer for efferent and afferent network connections to and from neuronal cortex layers. The Extended Data section provides figures and tables with information on mapped *m/z* values and a UMAP diagram highlighting value distributions for all categories.

**Fig. 4.**
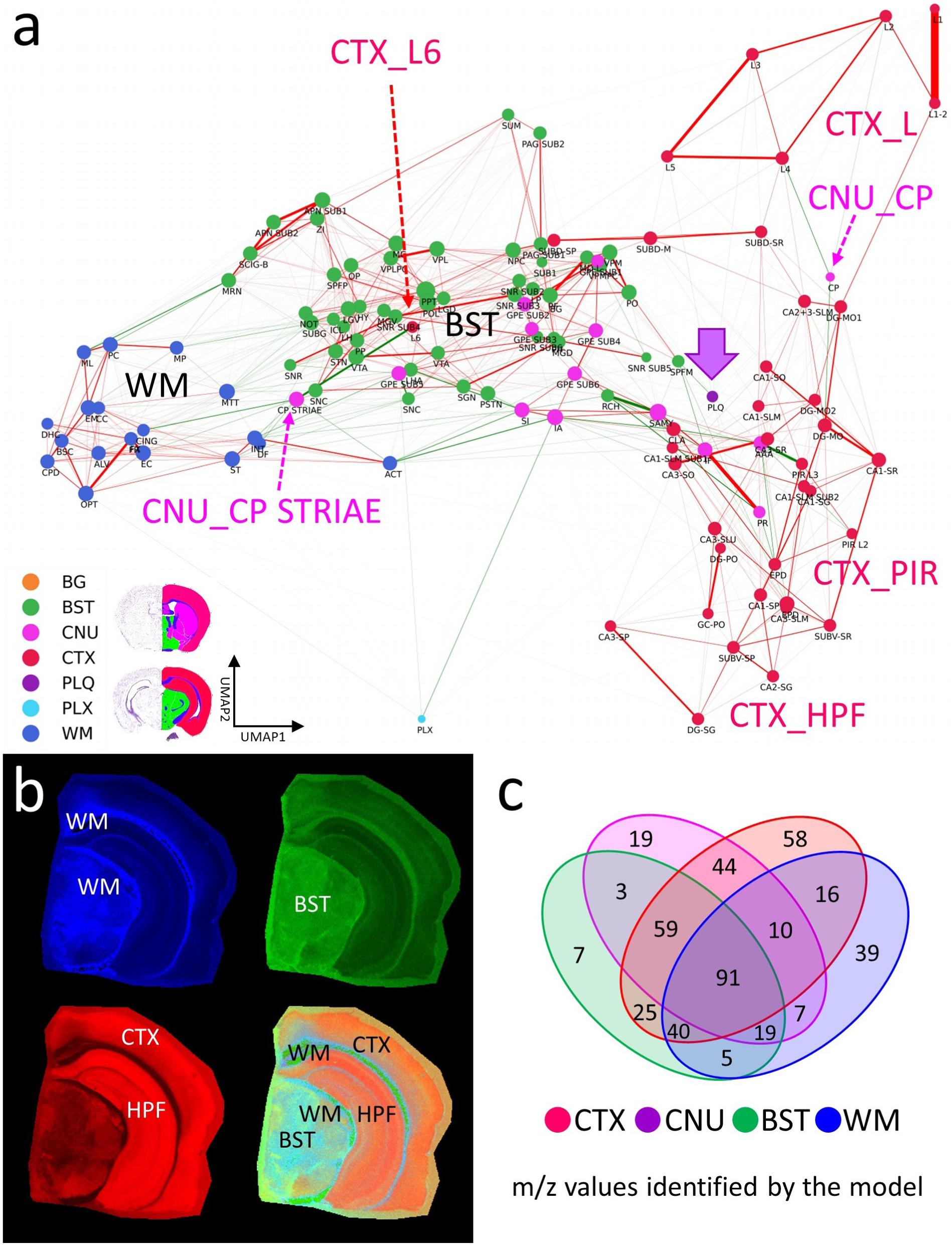
Brain regions have unique lipidomics profiles. **a**, The CBLA elucidates clustered distributions of major categories (color-coded) and clustered subcategories within the brain. Notably, the category corresponding to isocortex layer 6 (CTX L6) is located amid brainstem (BST) structures, attributable to its mixed composition of myelinated tracts and neuronal processes. Furthermore, caudate putamen striae (CNU CP STRIAE) within the cerebral nuclei (CNU) category signify myelinated tracts positioned between neurons of the caudate putamen (CNU CP) in the tissue. In the CBLA, these structures are situated between BST and white matter (WM). Also, the A*β* plaques category (violet arrow) is found among cortex (CTX) subcategories (CTX L, CTX PIR, and CTX HPF). **b**, RGB images visualize three major brain regions by summing mapped *m/z* values and removing shared values: blue, WM connections encompassing corpus callosum and brainstem WM; green, BST nuclei and connections; red, CTX, specifically isocortex layers (CTX L) and hippocampal formation (CTX HPF). **c**, Venn diagram highlighting *m/z* values identified for gross anatomical regions determined with ML models. Legend: BG, background; BST, brainstem; CNU, cerebral nuclei; CTX, cerebral cortex; PLQ, A*β* plaques; PLX, choroid plexus; WM, white matter.

### 3.4 Pathological Structures Reveal Origins and Effects on Brain Networks

A*β* plaques are small, dense, spherical deposits of aggregated A*β* protein. These toxic accumulations disrupt nearby neurons and their connections in the brains of patients and mouse models of Alzheimer’s disease. Using the CBLA, we explored the lipid composition and molecular relationships of these circumscribed pathological structures. During the annotation process, we generated more than 50 polygonal annotations grouped into the PLQ category, representing A*β* plaques.

The PLQ category shares numerous *m/z* values from distinct brain regions, indicating that plaque lipid signatures reflect contributions from their regions of origin (Fig. 5a,b). Using VPS of 1179.7308 *m/z* (GM3), plaques can be anatomically visualized in CTX L and PIR (Fig. 5c). In addition, we detected *m/z* features associated with disrupted neuronal connections, including dendritic and axonal networks (Fig. 6), demonstrating how CBLA enables unsupervised molecular phenotyping of pathological structures.

**Fig. 5.**
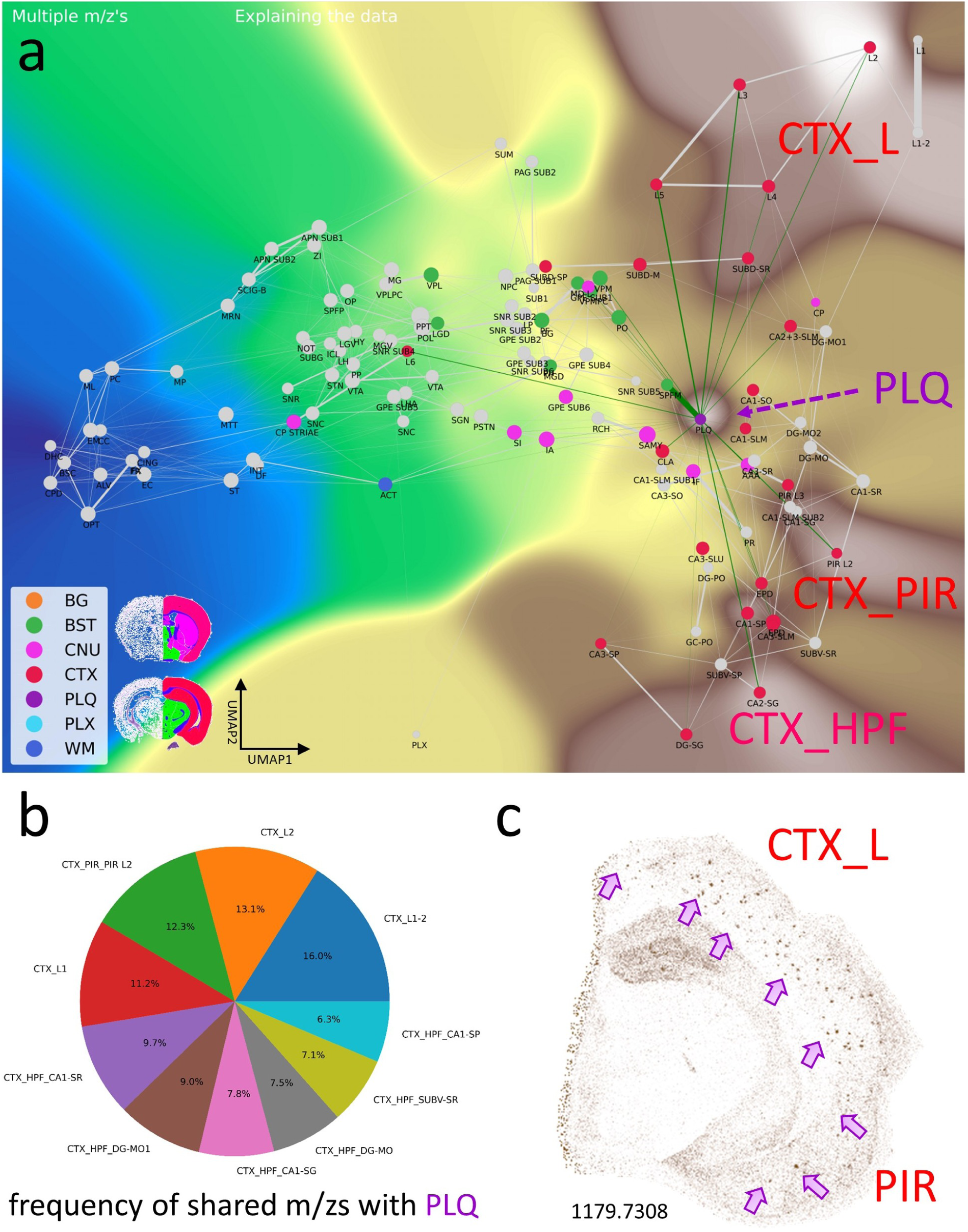
A*β* plaque-associated *m/z* signatures reveal relationships with several brain regions. The PLQ category exhibits an informative pattern of *m/z* values. This category has connections to various brain regions and is therefore located near the center of cortical brain regions: i) it comprises plaque-specific *m/z* values, and ii) it shares *m/z* values from different locations, e.g., CTX L (cortical layers), CTX PIR (piriform cortex), and CTX HPF (hippocampal formation). The PLQ node has edges to different brain regions; these are potential brain connections and indicate shared *m/z* values. Of note is the edge toward cortex layer 6 (CTX L6). **a**, *Virtual Landscape Visualization* (VLV) of *m/z* values significantly associated with the PLQ category. Prominent expression is observed in isocortex layers 2–5 (CTX L), piriform cortex (CTX PIR), and hippocampal formation (CTX HPF). **b**, Pie chart showing the top anatomical regions based on the frequency of *m/z* values shared with PLQ, predominantly within the isocortex (L) and hippocampal formation (HPF) of the CTX categories. **c**, A Virtual Pathology Stain (VPS) of *m/z* 1179.7308 reveals A*β* plaques in different isocortex layers (CTX L) and piriform cortex (PIR). Arrows indicate examples of individual plaques. Legend: BG, background; BST, brainstem; CNU, cerebral nuclei; CTX, cerebral cortex; PLQ, A*β* plaques; PLX, choroid plexus; WM, white matter; pie-chart labels follow Allen Atlas nomenclature.

**Fig. 6.**
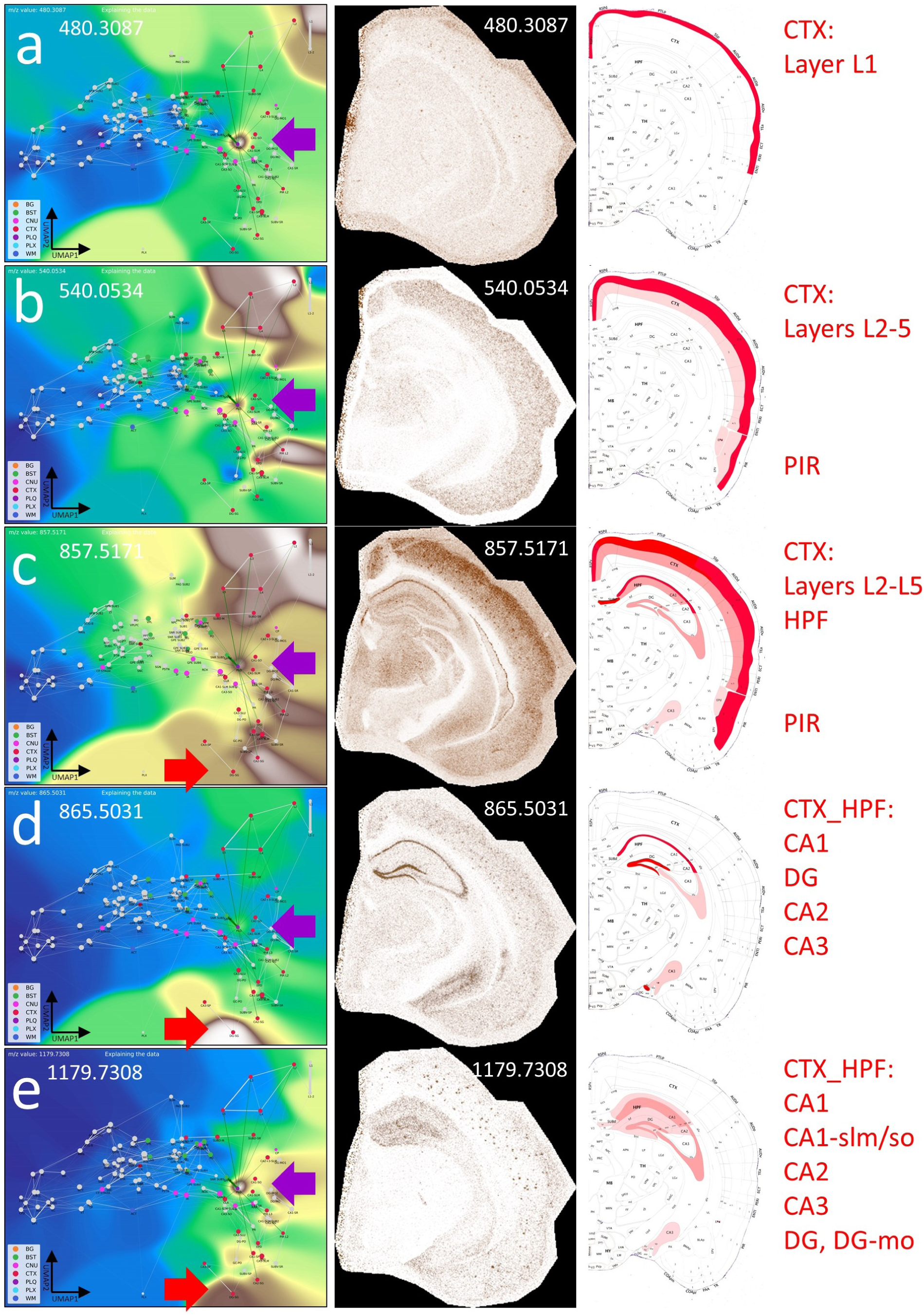
The CBLA maps A*β* plaque-associated lipid accumulation and brain connectivity. **a–e**, show the VLV for different *m/z* values associated with the PLQ category (violet arrow), their Virtual Pathology Stain (VPS) highlighting the corresponding brain regions, and a schematic annotation overlaid on the Allen mouse brain atlas [9], explaining the affected regions. In the VLV, only nodes that share PLQ *m/z* values are highlighted. A*β* plaques disrupt neural connections in the cortex (CTX L, CTX PIR) and accumulate lipids from degenerated axons. These lipid accumulations indicate, for example, the origin of disrupted connections and cause plaques to share *m/z* values with originating regions, e.g., from the hippocampus (CTX HPF, red arrow in VLV). Legend: BG, background; BST, brainstem; CNU, cerebral nuclei; CTX, cerebral cortex; PLQ, A*β* plaques; PLX, choroid plexus; WM, white matter.

By zooming in on individual plaques, we can infer their origin based on molecular patterns and spatial localization within the CBLA (Fig. 7). Using the VLVs, we further identified and validated low-abundance *m/z* m/z values enriched in A*β* plaques, providing insight into disrupted neuronal networks. Figure 8 illustrates an example of such a low-abundance m/z value, which is also detected in the hippocampal formation (CTX HPF), the supramammillary nucleus (SUM), and the choroid plexus (PLX).

**Fig. 7.**
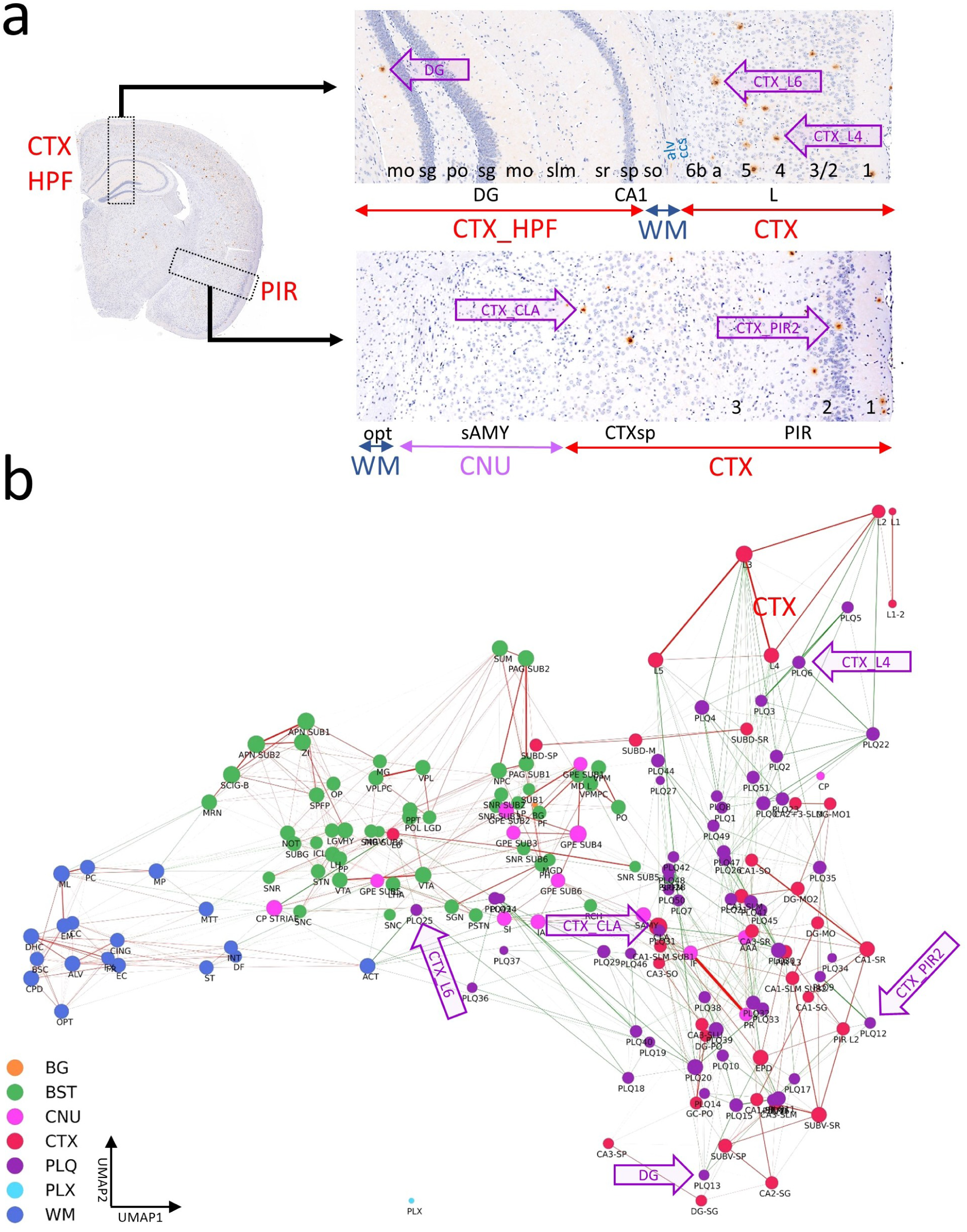
The CBLA uncovers location-specific lipid signatures of A*β* plaques. **a**, shows a brain hemisphere immunostained for A*β* and hematoxylin as counterstain, with two magnified views of specific CTX regions (boxed). The magnifications show the anatomical arrangement into specific CTX layers or HPF structures (red, CTX; blue, WM; violet, CNU). The annotations used to establish the CBLA are shown as text (mo*→*1 and 3*→*1). Arrows indicate examples of five plaques in different CTX regions (DG, L4, L6, PIR2, and CLA). **b**, In the CBLA graph, nodes corresponding to the labeled plaques (from **a**) are found near nodes of their originating anatomical locations, confirming their region-specific lipid content. Legend: BG, background; BST, brainstem; CNU, cerebral nuclei; CTX, cerebral cortex; PLQ, A*β* plaques; PLX, choroid plexus; WM, white matter.

**Fig. 8.**
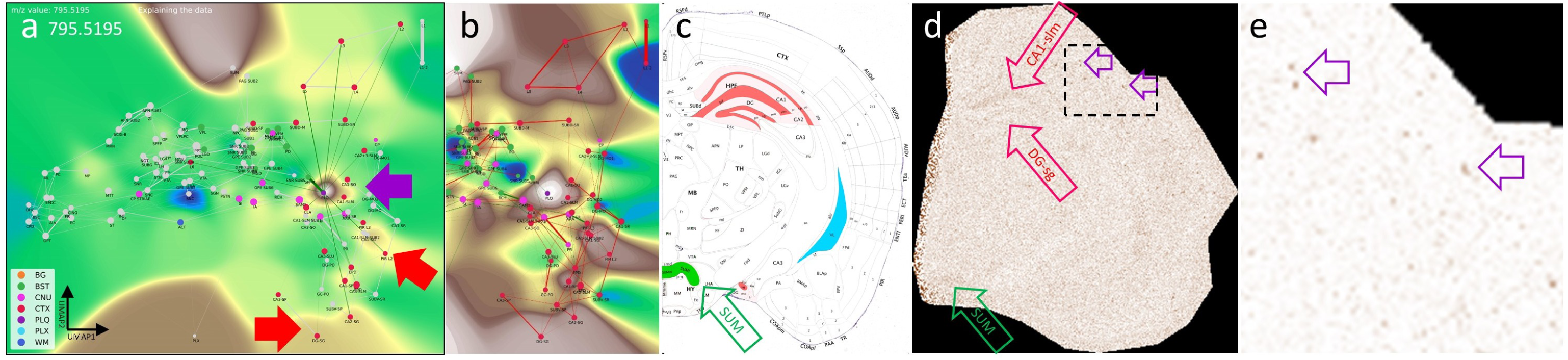
Visualizing a low-abundance A*β* plaque-related mass and its spatial relationship with other brain regions. **a**, The VLV of *m/z* 795.5195 shows a strong signal in the PLQ (violet arrow) and BST SUM categories, with lower intensity in PLX and CTX HPF DG-SG categories. **b**, Histogram-normalized VLV to increase visibility of low-abundance signals. **c**, Anatomical representation of *m/z* 795.5195 expression, color-coded by gross anatomical regions (see also Fig. 9). **d**, Linear VPS showing the *m/z* signal slightly above noise levels; anatomical regions are marked in correspondence with Panel c. Examples of A*β* plaques are indicated with violet open arrows. **e**, Magnification of Panel d and logarithmic VPS enhancing high-intensity signals and suppressing noise. A*β* plaques are indicated with violet open arrows and become clearly visible. A putative identification of the *m/z* values is given in Table S3. Legend: BG, background; BST, brainstem; CNU, cerebral nuclei; CTX, cerebral cortex; PLQ, A*β* plaques; PLX, choroid plexus; WM, white matter.

Interestingly, the distribution of these *m/z* values follows a molecular-weight-dependent pattern (Fig. 9), and each selected *m/z* value shows a putative anatomical origin based on its reproducible enrichment in specific subregions. This pattern suggests that plaque chemistry may integrate molecular inputs from defined, biologically connected brain regions rather than reflect random local accumulation; putative annotations for selected *m/z* values are provided in Table S3.

**Fig. 9.**
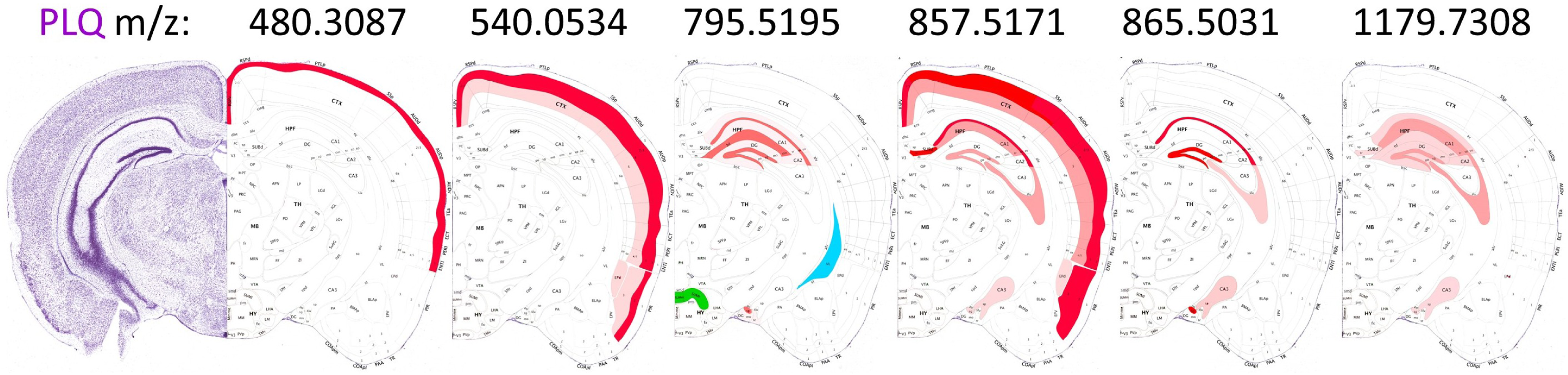
Schematic localization of six selected *m/z* features found in the PLQ category. The scheme highlights connections that contribute to the lipid profile of A*β* plaques, mostly originating from the isocortex (CTX L, CTX PIR categories) and the hippocampal formation (CTX HPF). The distribution changes with increasing *m/z*. The newly discovered *m/z* value 795.5195 is located in the supramammillary nucleus (BST SUM), which projects to CTX HPF, and in the choroid plexus (PLX). The Nissl stain and brain layout were modified from the Allen mouse brain atlas [9]. A putative identification of the *m/z* values is given in Table S3.

### 3.5 Generating New Biological Hypotheses with the CBLA

Our objective was to evaluate the CBLA framework for elucidating lipid distributions across functional brain networks. While mapping A*β* plaque *m/z* values and visualizing them using the CBLA and VLVs (Figs. 5-7), we observed a distribution pattern that correlated with plaque location of origin and its neuronal connections, such as hippocampal origin for *m/z* 1179.7308 (Fig. 6e). This led us to hypothesize that distinct lipid distribution patterns may be associated with specific functional-anatomical networks or individual neuronal connections in the brain. We therefore termed these lipids *index lipids*. Such patterns may reveal functional networks and provide novel insights into intricate lipid connectivity and biological interactions within the brain.

#### 3.5.1 Anatomical Regions and Functional Networks Show Specific Lipid Patterns

##### Nuclei and connections within the extrapyramidal motor network exhibit characteristic index lipids

Using the CBLA, we identified lipid distribution patterns that correlate with specific functional networks. Fig. 10 illustrates two examples of *m/z* values that reveal a shared network pattern consisting of nuclei and cortical regions involved in motor-function modulation. These lipid patterns could be used to investigate why neurodegenerative diseases have predilection sites at typical anatomical locations, how these diseases begin, and which factors promote pathogenic processes.

**Fig. 10.**
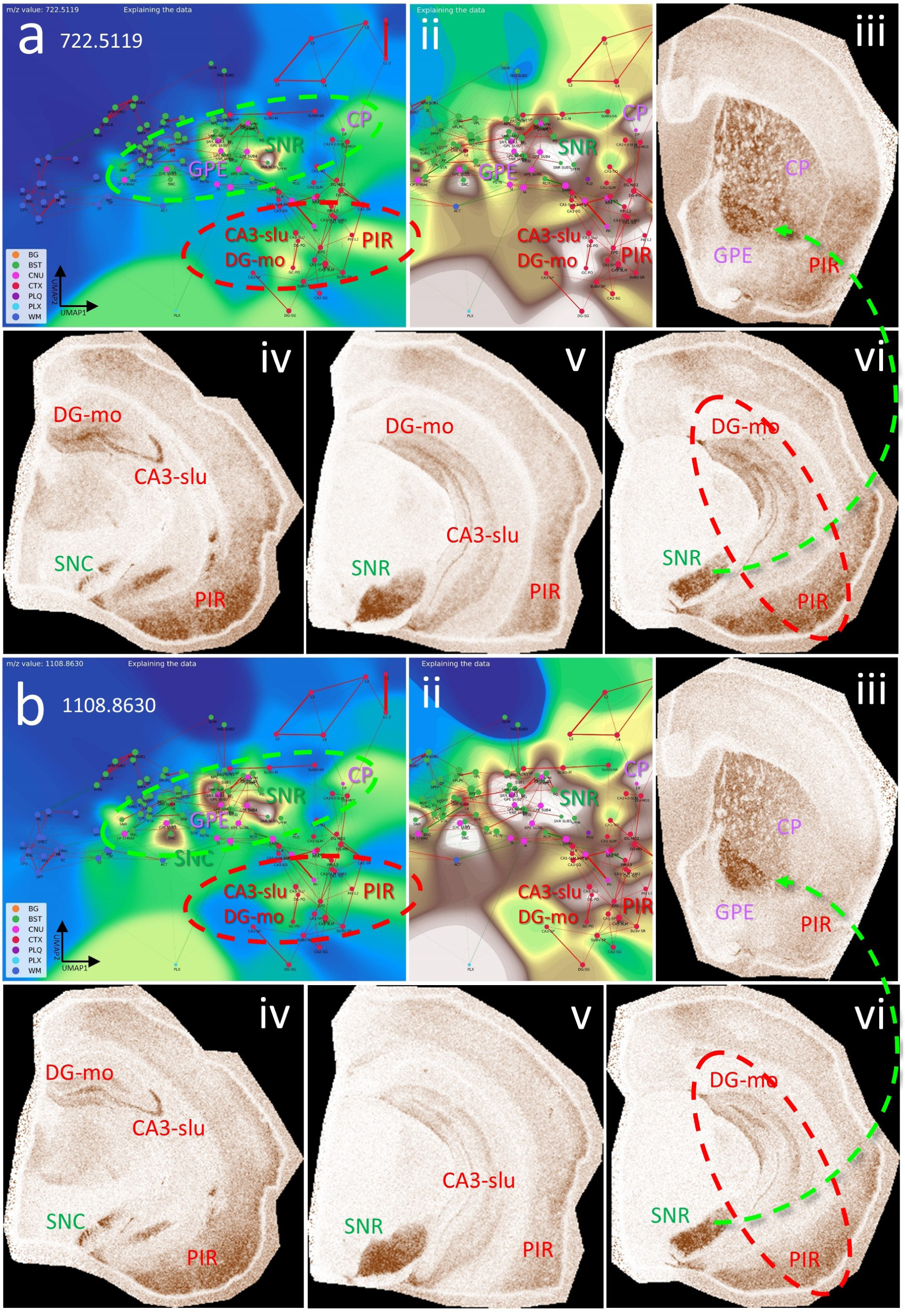
Nodes and connections of the extrapyramidal network share specific index lipids. **a**, CBLA for *m/z* 722.5119, highlighting its distribution in BST nuclei (green ellipse: SNR, SNC, and GPE), cerebral nuclei (CP), and cortical structures (red ellipse: CA3, PIR). Panel ii shows a histogram-normalized landscape to increase node visibility. Panels ii–vi show VPS views of *m/z* expression and distribution in SNC, SNR, DG-mo, CA3-slu, GPE, CP, and PIR. SNR/SNC are functionally connected to GPE/CP (iii and vi, green arrow), and PIR and HPF (DG, CA3) are also connected (vi, red ellipse). **b**, *m/z* 1108.8630 shows a nearly identical expression and distribution pattern to **a**, except for much higher expression in GPE (iii) and SNC (iv). Putative identifications of the *m/z* values are given in Table S3. Legend: BG, background; BST, brainstem; CA3, cornu ammonis part 3; CNU, cerebral nuclei; CP, caudate putamen; CTX, cerebral cortex; DG, dentate gyrus; GPE, globus pallidus, pars externa; HPF, hippocampal formation; PIR, piriform cortex; PLQ, A*β* plaques; PLX, choroid plexus; SNC, substantia nigra, pars compacta; SNR, substantia nigra, pars reticulata; WM, white matter. Panels iii–vi are shown at different fronto-occipital levels relative to bregma [14].

##### The hippocampal formation reveals organized lipidomic structures

The HPF consists of multiple functional-anatomical layers (see also Fig. 7a) that are connected to several cortical brain areas. These connections are affected during disease development and persistence, for example in temporal lobe epilepsy. Specific metabolic and lipidomic markers linked to these connections can be assessed to better understand the generation of grand mal seizures that originate locally in the HPF and their long-term effects on hippocampal-cortical network organization (Fig. 11).

**Fig. 11.**
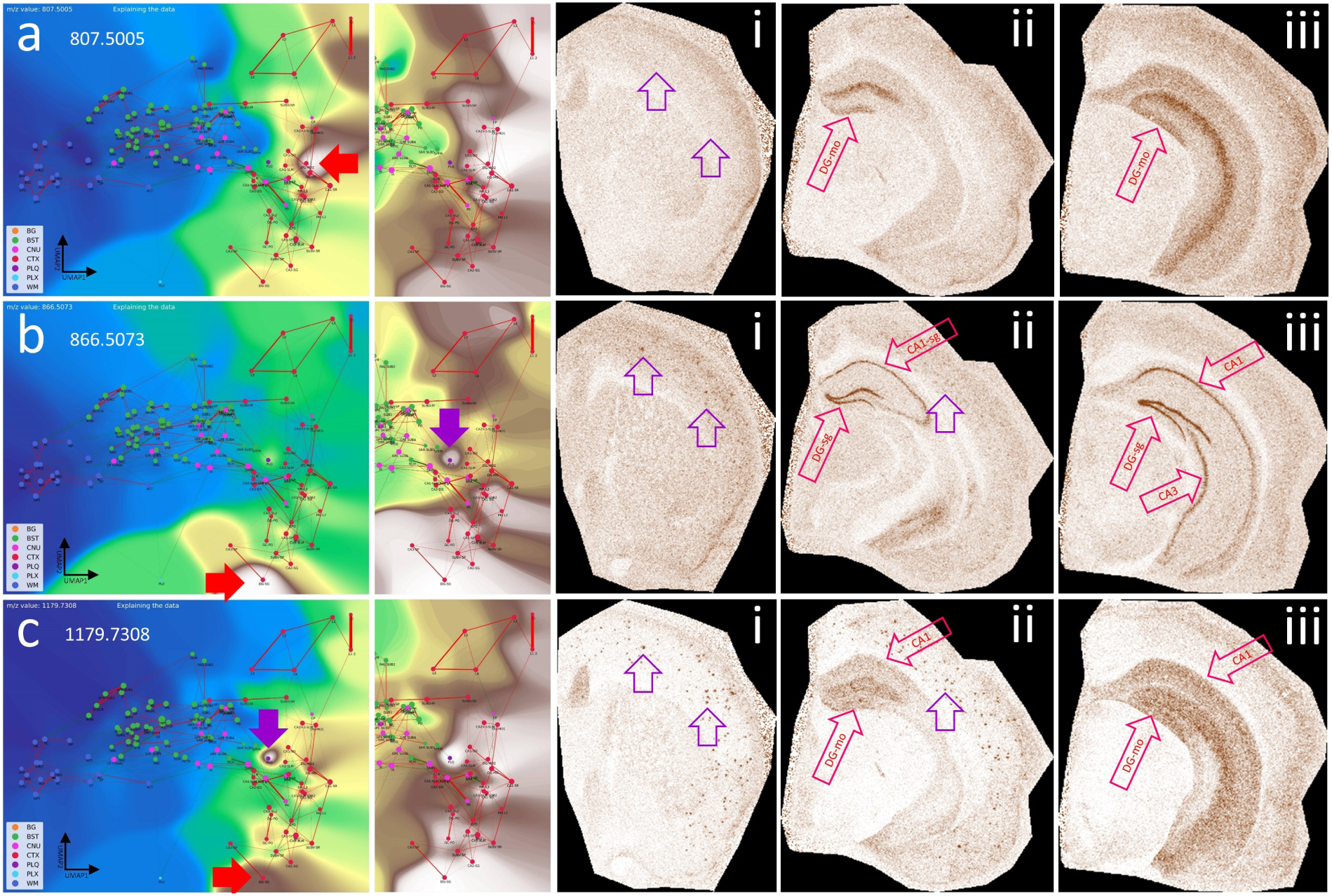
The hippocampal formation (HPF) has specific lipid signatures for its substructures. **a**, CBLA and a histogram-normalized view with the VLV for *m/z* 807.5005, which strongly emphasizes its specific localization in the molecular layer of the dentate gyrus (red arrows, DG-mo). Violet arrows indicate tiny speckles of A*β* plaques. **b**, characteristics of *m/z* 866.5073, showing the neuronal band of DG and CA1–3 as well as cortical A*β* plaques (i), revealing cortical connections of DG and CA. **c**, characteristics of *m/z* 1179.7308, showing extensive expression in connecting structures or neurons shown in **b**, and cortical A*β* plaques (i), again emphasizing disruption of hippocampal-cortical connections due to A*β* plaque deposition. Putative identifications of the *m/z* values are given in Table S3. Panels i–iii are shown at different fronto-occipital levels relative to bregma [14].

#### 3.5.2 Brainstem Nuclei Have Unique Lipid Compositions

The brainstem (BST) consists of diverse nuclei and intricate connections. We therefore hypothesized that BST lipid composition would show similarities to other brain regions. As depicted in Fig. 12, CBLA and VLV identified masses such as *m/z* 786.5286 and 881.5165 that are heterogeneously enriched within the BST, with certain nuclei, such as SUM (supramammillary nucleus) and APN (anterior pretectal nucleus), showing pronounced expression patterns. For these *m/z* values, some BST-to-cortex connections become visible, especially toward cortical layers 2 and 3. More broadly, knowledge of specific lipid patterns in BST nuclei may improve our understanding of complex neurodegenerative diseases. Several pathological processes in these diseases begin in specific brain ganglia or regions but are linked to the same pathological protein. One example is *α*-synuclein, which is found in several neurodegenerative diseases with motor impairment, including Parkinson’s disease, multiple system atrophy (MSA), and Lewy body dementia (LBD).

**Fig. 12.**
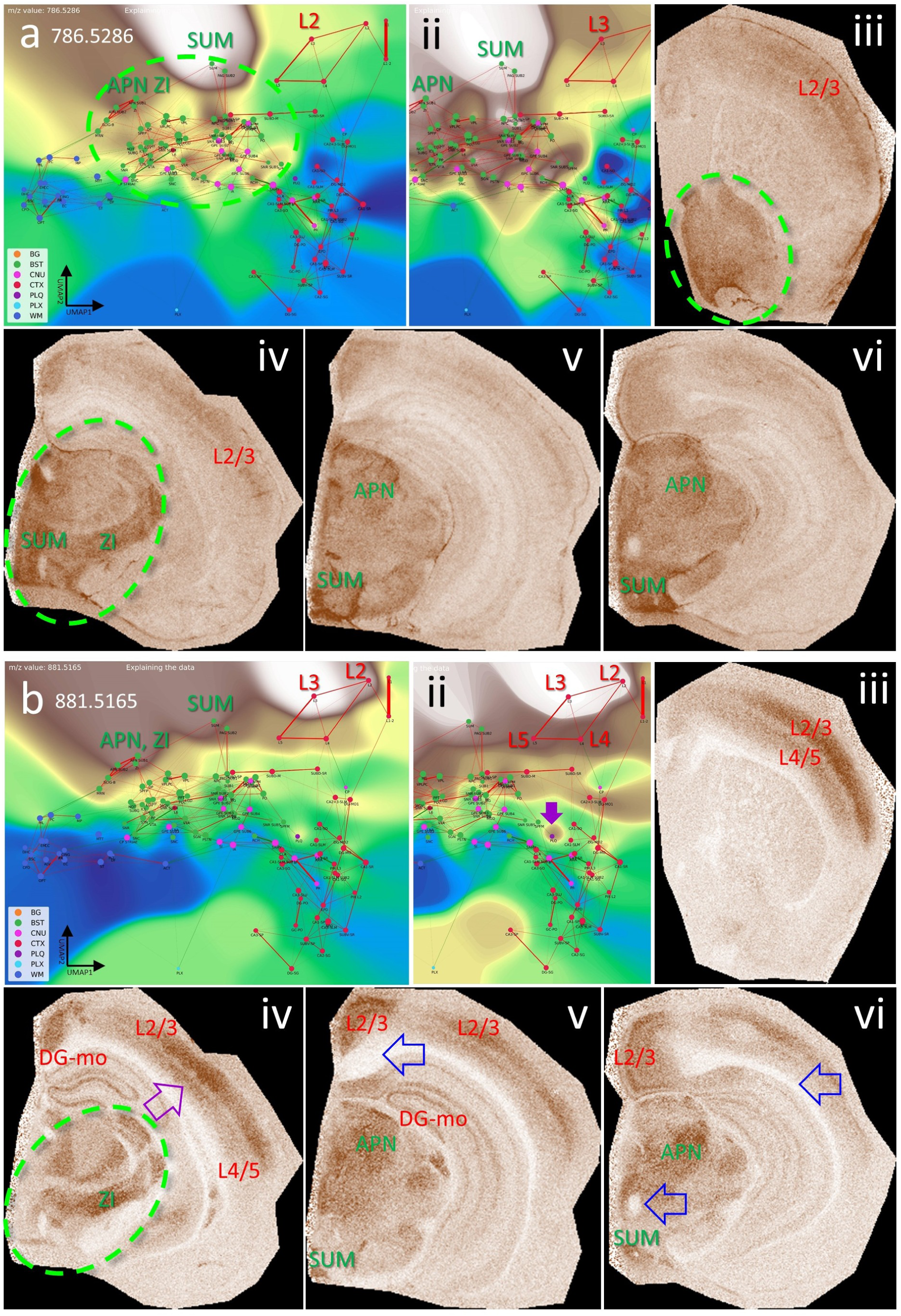
The brainstem (BST) has specific lipid signatures in its substructures and connections. **a, b**, show the CBLA and a histogram-normalized selection (ii) with VLV overlays of *m/z* 786.5286 and 881.5165, strongly emphasizing their localization in BST nuclei (green ellipse). **ii–vi**, show VPS at four locations in the mouse brainstem (green ellipse), highlighting specific nuclei. **a iv**, APN (anterior pretectal nucleus), SUM (supramammillary nucleus), and ZI (zona incerta) are examples of regions that exhibit higher expression than surrounding nuclei and tracts. **iii–vi**, cortex layers 2/3 and 4/5 (CTX L) show increased expression and represent input cortical layers for tracts from brainstem nuclei. **b iv–vi**, *m/z* 881.5165 also shows specific expression in HPF regions, specifically in DG-mo and CA1-sr/slm. **b v–vi**, open blue arrows point to WM tracts that are fully negative, and in **b ii + iv**, violet arrows point to tiny A*β* plaque residues from destroyed cortical projections. Legend: BG, background; BST, brainstem; CNU, cerebral nuclei; CTX, cerebral cortex; PLQ, A*β* plaques; PLX, choroid plexus; WM, white matter. Panels iii–vi are shown at different fronto-occipital levels relative to bregma [14].

#### 3.5.3 White Matter Tracts Show Distinct Lipid Content

White matter (WM) contains a large proportion of brain lipids and provides effective insulation of neuronal connections, thereby increasing conduction speed. WM establishes connections between different brain regions, for example between the two hemispheres through the corpus callosum (cc), and also forms long intrahemispheric tracts extending from frontal to occipital lobes. Using CBLA and VLV, we identified lipid signatures specific to WM (see 6), exemplified by *m/z* 756.5894 in Fig. 13. Thus, the combination of CBLA and VLV enables identification of comprehensive lipid profiles that are shared across, or unique to, specific anatomical brain regions and can serve as a resource for generating novel biological hypotheses. Moreover, these tools can be expanded with other MSI modalities, such as metabolomics and proteomics, to integrate complex region-specific information patterns.

**Fig. 13.**
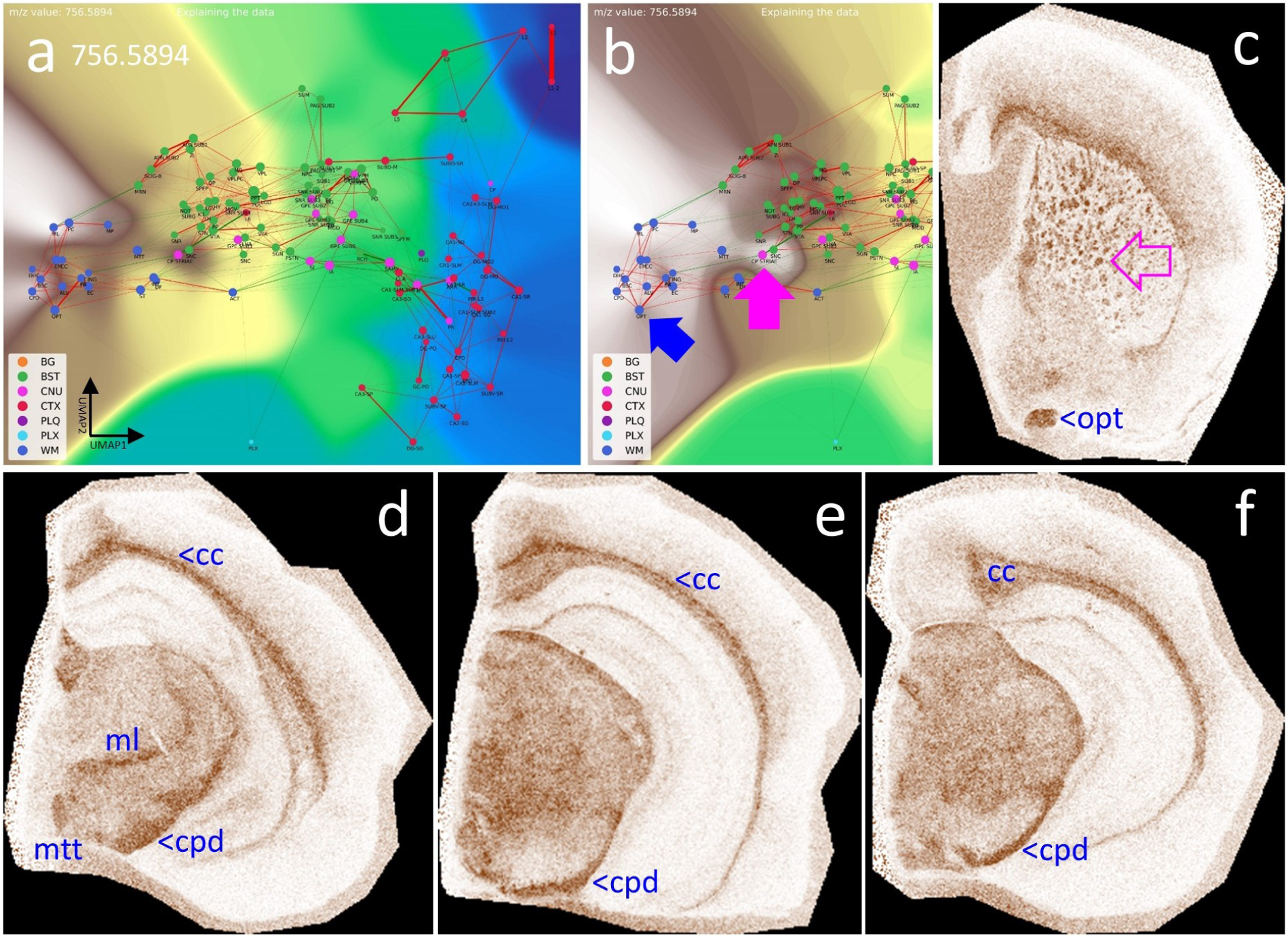
White matter tracts (WM) have specific lipid signatures. **a**, CBLA and a histogram-normalized selection (**b**) with a VLV overlay for *m/z* 756.5894, which strongly emphasizes its specific localization in WM tracts (left cluster, dark blue nodes) and lower abundance in BST tracts (green nodes). The blue arrow points to the optic tract (opt) as a representative example highlighted in (**c**); the pink arrow points to the striae in the caudate putamen (CP striae), which are numerous small white matter tracts (**c**) crossing the CP from BST nuclei to the CTX. **c–f**, show four VPS at different locations to visualize the in situ distribution in WM tracts in the cc and BST. Legend: BG, background; BST, brainstem; CNU, cerebral nuclei; CTX, cerebral cortex; PLQ, A*β* plaques; PLX, choroid plexus; WM, white matter; cc, corpus callosum; cpd, cerebral peduncle; ml, medial lemniscus; mtt, mammillothalamic tract; opd, optic tract. Panels c–f are shown at different fronto-occipital levels relative to bregma [14].

### 3.6 Using the CBLA to Visualize Region-Specific Differences Between Mouse Models

Finally, we aimed to demonstrate the applicability of the CBLA to general research questions, including studies of disease mouse models. In our dataset, we analyzed tissue from a new mouse model with deficiency of the ABCA7 lipid transporter [13]. ABCA7 is a major genetic risk factor for Alzheimer’s disease and mediates lipid transport across the plasma membrane and into the intracellular space (cytoplasm) [32, 33]. Therefore, investigating its biological function using MSI and appropriate mouse models is of high interest in neurodegenerative disease research. Using CBLA, we show for the first time examples of MSI-derived *m/z* values that are differentially expressed between ABCA7-deficient mice and non-deficient control mice (Fig. 14). These *m/z* values represent lipids linked to ABCA7-dependent distribution patterns in the brain and may reflect disease-modifying lipids or lipids directly involved in pathogenic processes.

**Fig. 14.**
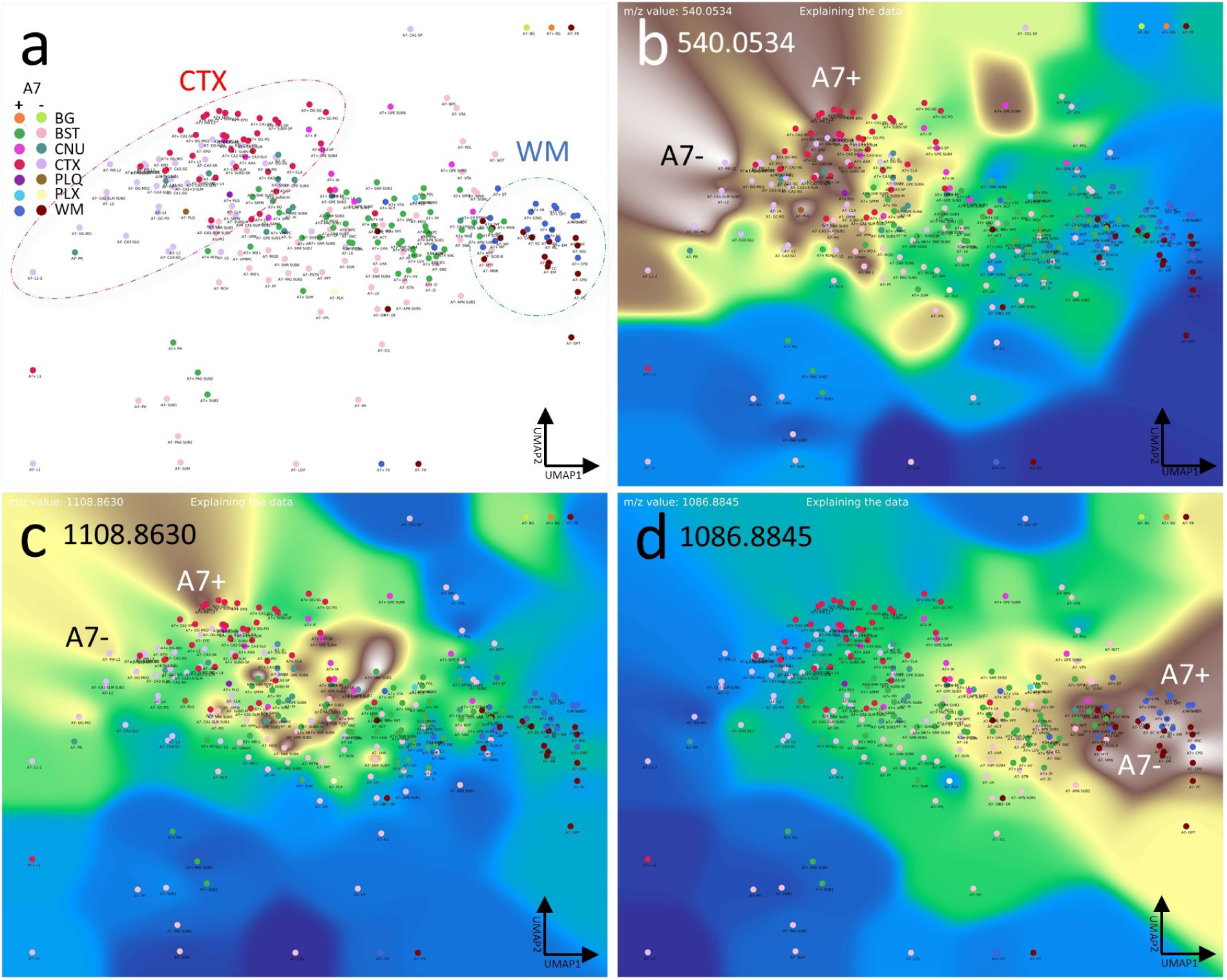
Lipid transporter ABCA7-deficiency results in altered brain lipid distribution and abundance profile. **a**, CBLA. Not all annotation categories appear in all mice; therefore, we designed an ML model to generate annotations for this comparison. We ran an ensemble of 100 models and used the majority prediction. We grouped the mice into A7–and A7+ cohorts. For the nodes, we measured the average spectra for each predicted brain region type without TIC normalization to enable comparison of raw intensity differences between mice. The circles highlight the CTX (red) and WM (blue) nodes, which are clustered but show a distinct shift in their location due to slight differences in their composition between the A7+ (wild-type) and the A7– (deficient) animals. **b–d**, show examples of three VLVs of differentially expressed *m/z* values, highlighting either A7– or A7+ preference. Panels b and d highlight *m/z* values increased in A7– mice, whereas Panel c shows an *m/z* reduced in A7– mice (increased in A7+ controls). Legend: BG, background; BST, brainstem; CNU, cerebral nuclei; CTX, cerebral cortex; PLQ, A*β* plaques; PLX, choroid plexus; WM, white matter

## 4 Discussion

This study presents the first large-scale brain annotation and mapping effort in mice based solely on Mass Spectrometry Imaging (MSI) data, covering a substantial portion of the categories defined in the Allen mouse brain atlas [9]. To manage the complexity of high-dimensional MSI data, the large number of anatomical regions, and their relationships to the spatial distribution of selected *m/z* features, we developed the *Computational Brain Lipid Atlas* (CBLA). CBLA improved anatomical annotation while enabling the generation of new biological insights and hypotheses. In contrast to previous studies that focused mainly on broad anatomical annotation [5, 6, 34], our approach captures fine brain substructures without requiring additional modalities such as immunostaining [6]. By integrating multiple continuous visualization panels [2] derived directly from raw MSI data, we streamline annotation and enable a substantially larger set of brain regions to be labeled.

Our findings show that MSI, even when applied to a relatively small lipidomics dataset with limited spatial resolution as in this study, can resolve detailed brain anatomy and serve as a foundational tool for biological exploration.

Using CBLA, we provide evidence that lipid distributions exhibit region-specific signatures, allowing brain subregions to be differentiated solely by lipid composition. This work also provides a comprehensive mapping of *m/z* values linked to specific brain regions and uses it to examine the spatial organization of A*β* plaques. Our analysis suggests that these plaques retain *m/z* signals from their region of origin and display spatial patterns that mirror functional connectivity, including links to distant brain areas. These findings arise from the combined *m/z* mapping and CBLA visualization framework. Beyond the present AD-focused examples, the same strategy is applicable to other neurodegenerative models in which pathology follows vulnerable networks (e.g., hippocampocortical, extrapyramidal, or brainstem-associated systems). In this context, MSI-ATLAS can support cross-model comparisons of region-resolved molecular signatures, disease-stage trajectories, and treatment-associated shifts in network chemistry.

These observations are consistent with recent MSI studies in AD-related tissue that report plaque heterogeneity, region-dependent lipid signatures, and isotope-informed spatial dynamics of plaque-associated chemistry [35–40].

Several limitations should be noted. First, this study is based on a relatively small cohort and a single acquisition configuration, which limits statistical power and generalizability across laboratories, platforms, and tissue-preparation workflows. Second, to resolve *m/z* values beyond fixed binning resolution, we developed a Gaussian mixture model (GMM)-based approach [21]. Although this improves precision, it introduces a trade-off: increased sensitivity to subtle differences can split a single biological signal into multiple closely related peaks. One strategy to mitigate this issue is to incorporate spatial priors, allowing *m/z* resolution within the anatomical context of annotated regions. Alternatively, Trapped Ion-Mobility Time-of-Flight (TIMS-TOF) mass spectrometry [41] may further disambiguate signals and reduce redundancy. A third challenge is the annotation of highly similar subregions. Our workflow addressed this through iterative refinement with complementary visualization panels, each emphasizing distinct structural features. Future improvements should include active-learning strategies that prioritize uncertain or under-annotated regions, explicit quantitative region-annotation confidence scores, and external validation on independent neurodegenerative cohorts. These steps would improve robustness, reproducibility, and translational utility.

In addition, we further note several workflow-level limitations. First, we used NEDC because it provides robust negative-ion lipid signals and stable spatial contrast in our acquisition setting; however, alternative matrices and complementary ionization conditions may broaden molecular coverage and should be explored in follow-up work. Second, model interpretation can be affected by isotope/adduct redundancy and by non-lipid or off-tissue/interferent ions, which may receive substantial weights if not explicitly filtered. Third, we used TIC normalization for a consistent global workflow, but internal-standard-based normalization is an important future extension for improving quantitative robustness across sections and batches. Finally, multimodal integration (e.g., MSI with microscopy/image-fusion pipelines) is a clear next step to strengthen structural interpretation and biochemical assignment while preserving the MSI-native annotation strategy.

Regarding biochemical interpretation, not all highlighted *m/z* features can currently be assigned with high confidence (Table S3). The table summarizes putative annotations for selected *m/z* features highlighted in the main text figures. Where feasible, we provide tentative assignments and treat them as hypothesis-generating rather than definitive identifications. In this context, the signal around *m/z* 1179.7308 is discussed in relation to prior reports of ganglioside-related species in AD models and patients [37, 42, 43], including GM3(36:1), and should therefore be regarded as a biologically plausible candidate pending orthogonal validation. This interpretation is supported by its detection in both hippocampal structures and cortical plaques, where lipid accumulation may reflect disrupted hippocampocortical connectivity [44, 45] (Fig. 5c, Fig. 6e, and Fig. 9). This observation raises the hypothesis that disruption of this pathway may contribute to cortical A*β* deposition patterns linked to a hippocampal origin.

## 5 Conclusions

MSI data are inherently rich and informative, yet remain underexploited because tools for high-resolution and interpretable analysis are still limited. The MSI-ATLAS presented here provides a large-scale, interpretable map of lipid distributions across brain regions and establishes a foundation for future studies of molecular brain architecture that bridge spatial metabolomics and neuroanatomical function.

## Supporting information

R1 response letter for Free Neuropathology

## 6 Declarations

### Ethics Approval

Animal breeding and tissue harvesting was approved by the local authorities (IV2-2022).

### Consent to Participate and Consent for Publication

Not applicable.

### Availability of Data and Material

The datasets generated and analyzed during the current study are available in the ProteomeXchange PRIDE repository under accession number PXD056609.

### Availability of Code

Code is available as an open-source package on [GitHub](https://github.com/jacobgil/msi-atlas)

### Competing Interests

The authors declare no competing interests.

### Funding

J.P. received funding from Nasjonalforeningen for folkehelse (Demensforskningsprisen 2025, Norway), Norges forskningsråd [NFR, Norway; 327571 ([PETABC](https://pahnkelab.eu/funding/petabc-jpnd/)), 295910 (NAPI), HelseSØ (Norway; 2022046), and the EIC Pathfinder Open Challenges program (European Commission; 101185769). The Proteomics Core Facility at the University of Oslo and Oslo University Hospital is supported by the Core Facilities Program of the South-Eastern Norway Regional Health Authority (HSØ) and NAPI (www.napi.uio.no, NFR, Norway; 295910).

### Authors’ Contributions

Conceptualization (J.G. and J.P.), Methodology (J.G. and J.P.), Software (J.G. and J.P.), Validation (J.G. and J.P.), Formal analysis (J.G., J.S., and J.P.), Investigation (J.G., J.S., and J.P.), Resources (J.P.), Data curation (J.P.), Writing - original draft (J.G. and J.P.), Writing - review and editing (J.G., J.S., and J.P.), Visualization (J.G. and J.P.), Supervision (J.P.), Project administration (J.P.), Funding acquisition (J.P.).

## Acknowledgements

We thank Thomas Brüning and Iván Eiriz for preparing the mice, brain samples, and Intellislides for MSI measurements.

## List of Abbreviations

Anatomical abbreviations were used according to the Allen mouse brain atlas nomenclature [9].

A*β* - amyloid-*β*, A7– - ABCA7-deficient mice [13], A7+ - ABCA7 non-deficient (wild-type) mice, APPtg - APP-transgene, APN - anterior pretectal nucleus, BG - background category, BST - brainstem category, CA2–sg - cornu ammonis area 2, stratum granulosum, CA3 - cornu ammonis area 3, CA3-slu - cornu ammonis area 3, stratum lucidum, CBLA - Computational Brain Lipidomics Atlas, cc - corpus callosum, CNU - cerebral nuclei category, CP - caudate putamen, cpd - cerebral peduncle, CTX - cortex category, CTX L - cerebral cortex layers, DG - dentate gyrus, DG–mo - dentate gyrus, molecular layer, DG–sg - dentate gyrus, stratum granulosum, EPD - endopiriform nucleus, dorsal part, GMM - Gaussian mixture model, GPE - globus pallidus, pars externa, HPF - hippocampal formation, IA - intercalated amygdaloid nuclei, IHC - immunohistochemistry, L1*/*2 - cortical layers 1/2, ML - machine learning, ml - medial lemniscus, mPAUC - *m/z*-Pairwise-AUC, MRI - magnetic resonance imaging, MS - mass spectrometry, MSI - mass spectrometry imaging, MSI–ATLAS - Mass spectrometry imaging atlas computational tool, MSI–VISUAL - Mass spectrometry imaging visualization tool [2], mtt - mammillothalamic tract, NEDC - N-(1-naphthyl) ethylenediamine dihydrochloride, opd - optic tract, PCA - principal component analysis, PIR - piriform cortex, PLQ - A*β* plaque category, PLX - plexus category, SNC - substantia nigra, pars compacta, SNR - substantia nigra, pars reticulata, SUM - supramammillary nucleus, TIC - total ion current, TIMSTOF - trapped ion mobility time-of-flight, UMAP - Uniform Manifold Approximation and Projection, VLV - Virtual Landscape Visualization, VPS - Virtual Pathology Stain, WM - white matter category, ZI - zona incerta.

## Supplementary Materials

Figures S1 to S3

Tables S1 to S3

### Comparison of *m/z* Selection Strategies for Plaque-Associated Signals

**Fig. S1.**
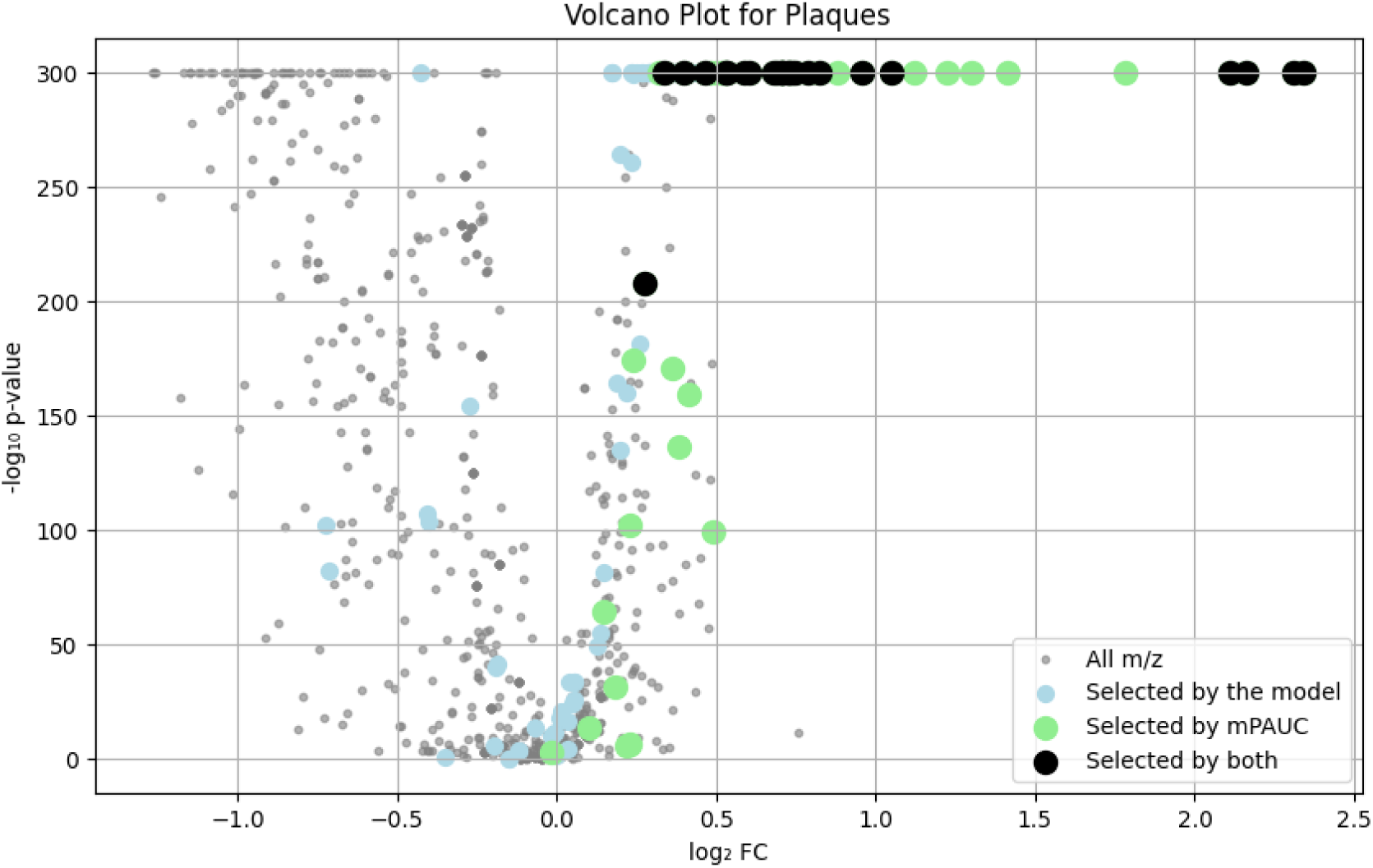
Different *m/z* selection strategies applied to the A*β* plaque (PLQ) category. The model-based approach also selects *m/z* values that are shared with other categories and can therefore have a negative log_2_ fold-change. The mPAUC method is selective for *m/z* values that are specific to PLQ because it compares values pairwise. The intersection of both methods results in highly selective masses for PLQ, while their union may help gather all potential *m/z* values associated with PLQ.

### Model-Explanation View of CBLA Category Weights

**Fig. S2.**
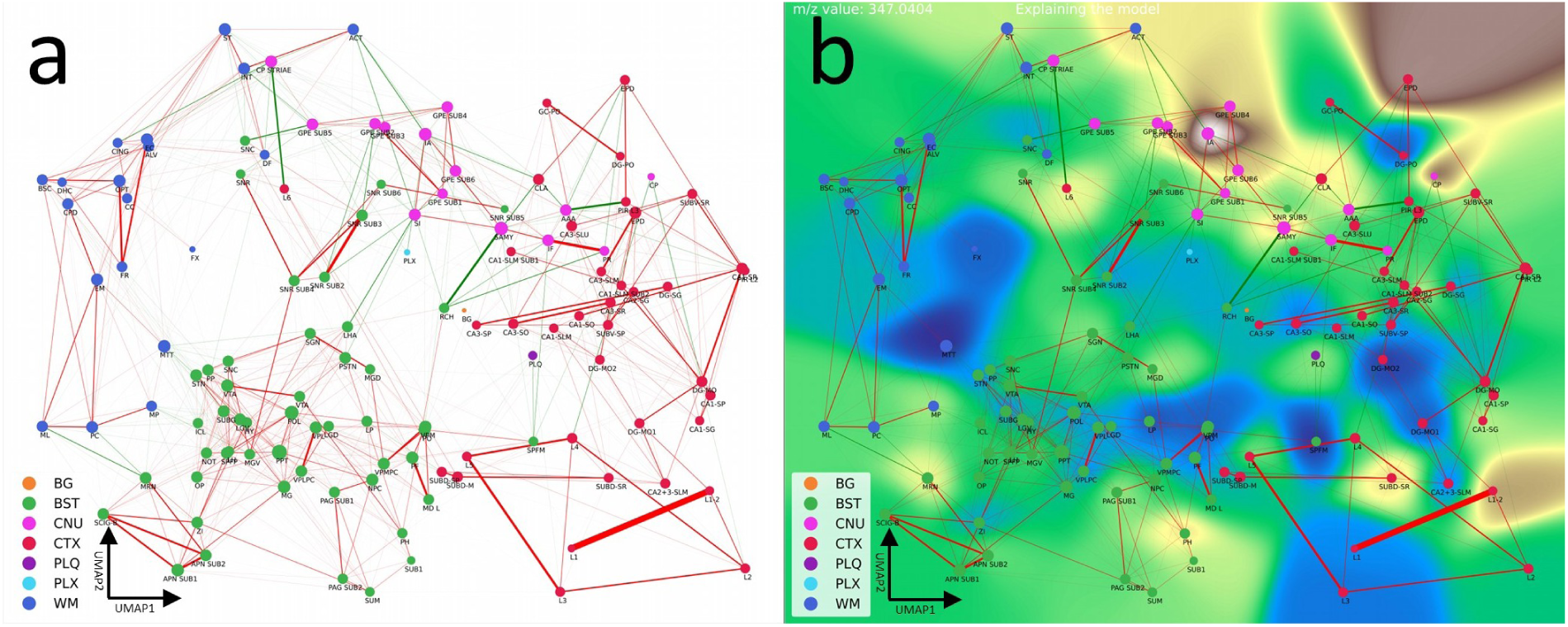
Explaining the model. **a**, Graph of the CBLA used for explaining the model weights, where we used a Bayesian logistic regression model for classifying the categories and using the category weights for each node before applying dimensionality reduction. **b**, VLV for explaining the model weights. Here, the nodes represent the learned model weights for each category. The model weight coefficients for the *m/z* 347.0404 are overlaid with the ‘terrain’ Matplotlib color scheme. Notice that, like for the data VLV (Fig. 4), the IA and EPD categories show a strong response, confirming the model has found this *m/z* predictive for these categories. However, unlike the data VLV, the model does not show a strong signal for the CA2-sg or the L1–2 categories, potentially showing that the model could be improved. Legend: BG, background; BST, brainstem; CNU, cerebral nuclei; CTX, cerebral cortex; PLQ, A*β* plaques; PLX, choroid plexus; WM, white matter.

### Comparing Lipidomic Profiles per Brain Region

#### *m/z* Mapping Data

The CSV files with mappings of *m/z* values to categories can be found here:

1. Table S1: model-based mapping (CSV file)
2. Table S2: mPAUC mapping (CSV file)

### UMAP Visualization of All Annotations

The UMAP visualizations of the *m/z* content of the annotations are given in the Extended Data, Fig. S3, showing that different brain regions have distinct lipidomic profiles, enabling their visualization and annotation.

**Fig. S3.**
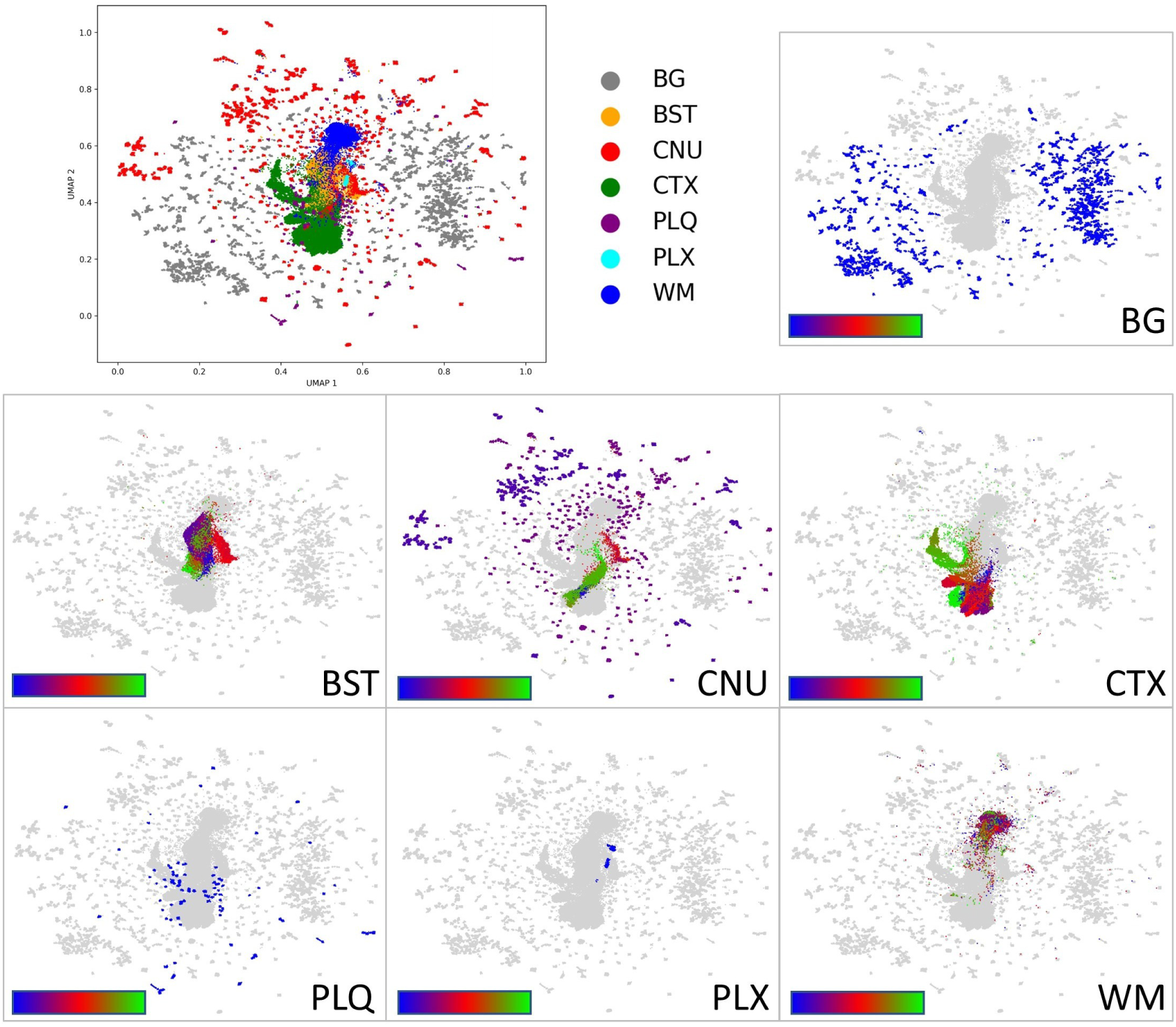
Analyzing lipid profiles of brain regions. The high-dimensional MSI data was reduced to two dimensions using UMAP [16]. The 123 regions were grouped into the main categories according to their anatomical locations [9] or designated as background (BG; ground truth). Each main category was also colorized with a BRG color scheme, showing the diversity within the categories. This visualization shows that annotated points from the same categories tend to distribute similarly in the low-dimensional visualization, which would be expected if the annotations were correct. On the other hand, our annotations capture extreme diversity in the data, especially in the CNU, WM, CTX, and BST categories. Legend: BG, background; BST, brainstem; CNU, cerebral nuclei; CTX, cerebral cortex; PLQ, A*β* plaques; PLX, choroid plexus; WM, white matter.

#### Identification of selected *m/z* features

Putative lipid annotations were assigned as described in Methods (see Putative lipid annotation), and are reported here as hypothesis-generating assignments requiring orthogonal validation (e.g., MS/MS).

**Table S3.** Putative identification and spatial localization of selected *m/z* features.

| $m/z$ | Putative annotation | Associated regions |
| --- | --- | --- |
| 347.0404 | IMP | CP, GPE, EPD, SAMY_IA, SAMY_AAA, CA2-sg, CLA |
| 480.3097 | LysoPE(18:0) | CTX.L1 |
| 540.0534 | cyclic ADP-ribose | CTX.L1–L5, PIR |
| 722.5119 | PE O-36:5 | SNR, SNC, GPE, CP, CA3, PIR |
| 795.5195 | PG 38:5 | SUM, PLQ, DG, CA1, CA2 |
| 807.5005 | PI(32:1) class | DG-mo, HIP |
| 857.5171 | PI(36:4) class | CTX.L1–L5, HIP, PIR |
| 865.5031 | PI/PG (unresolved) | CA1, DG, CA2, CA3 |
| 1179.7308 | GM3(36:1) | PLQ, CA1, CA2-slm/so, CA2, CA3, DG, DG-mo |

## Abbreviations

AAA: anterior amygdalar area
ADP: adenosine diphosphate
CA1: cornu ammonis area 1
CA2: cornu ammonis area 2
CA2-sg: cornu ammonis area 2, stratum granulosum
CA2-slm/so: cornu ammonis area 2, stratum lacunosum-moleculare/stratum oriens
CA3: cornu ammonis area 3
CLA: claustrum
CP: caudate putamen
CTX L1: cerebral cortex layer 1
CTX L1–L5: cerebral cortex layers 1 to 5
DG: dentate gyrus
DG-mo: dentate gyrus, molecular layer
EPD: endopiriform nucleus, dorsal part
GM3: monosialodihexosylganglioside (ganglioside GM3)
GPE: globus pallidus, pars externa
HIP: hippocampus
IA: intercalated amygdaloid nuclei
IMP: inosine monophosphate
PE: phosphatidylethanolamine
PG: phosphatidylglycerol
PI: phosphatidylinositol
PIR: piriform cortex
PLQ: A*β* plaque category
SAMY: striatum-like amygdalar nuclei
SNC: substantia nigra, pars compacta
SNR: substantia nigra, pars reticulata
SUM: supramammillary nucleus

## Notes

https://github.com/jacobgil/msi-atlas

### Competing Interest Statement

The authors have declared no competing interest.

### Summary of Updates

R1 version for Free Neuropathology / substantial text and figure changes

