## Supplementary material for "Mass spectrometry imaging-based explainable machine learning reveals the biochemical landscapes of the mouse brain": R1 response letter for Free Neuropathology

### Response to Editor and Reviewers (Round 1)

#### 1. Letter to the Editor

Dear Werner,

Thank you for the opportunity to revise our manuscript and for coordinating the peer-review process at *Free Neuropathology*. We are grateful to both reviewers for their constructive and insightful comments.

We have carefully revised the manuscript in response to all comments. The revised submission package includes:

- a clean revised manuscript;
- a marked manuscript with tracked/visible changes; and
- this point-by-point response document.

In summary, we clarified wording and terminology throughout the manuscript, improved figure legends and notation consistency, expanded the Discussion to address methodological limitations and future directions, incorporated the requested references, added putative lipid identifications for highlighted *m/z* features (with expanded context for plaque-associated *m/z* 1179.7308), and streamlined selected figure content to improve readability.

In addition, we performed a manuscript-wide language and consistency edit to improve grammar, phrasing, and terminological harmonization while preserving the scientific content and conclusions.

A concise list of major revisions is provided below, followed by detailed responses to each reviewer comment.

We believe these revisions have substantially improved the clarity, rigor, and readability of the manuscript, and we hope the revised version is suitable for publication.

Sincerely,

Jens Pahnke, Jacob Gildenblat, and Jorunn Stamnæs

#### 2. List of Major Changes

1. Clarified terminology and wording in title, abstract, introduction, and results to improve precision and reduce potential overstatement.

2. Expanded figure legends and formatting consistency (including abbreviation definitions and notation consistency for *m/z*).

3. Added discussion of workflow limitations, including potential biases (isotopes, adducts, off-tissue/interferent ions, and normalization considerations).

4. Expanded contextual discussion on future methodological extensions (additional ionization matrices and multimodal integration, including image-fusion directions).
5. Added citations requested by Reviewer A (PMID: 37006756, 36856542, 34882796, 38299453, 40274785, 40890115).
6. Added putative lipid identifications for highlighted *m/z* features in supplementary material and expanded biological context for plaque-associated *m/z* 1179.7308 (including ganglioside-related interpretation).
7. Revised figure captions and supplementary visualization annotations to improve section provenance clarity (including serial-section context) and delineation of on-/off-tissue signal interpretation.
8. Revised organization and readability, including streamlining/relocating non-essential figure content to supplementary materials where appropriate.
9. Updated figure numbering after relocation to supplementary materials: former Figure 10 is now Supplementary Figure S1, and former Figure 13 is now Supplementary Figure S2.
10. Corrected grammar/style issues, clarified TIMS/ion-mobility usage in Methods, standardized *m/z* terminology, and removed duplicated conclusion text in the Discussion section.
11. Performed an additional sentence-level language polish in the Introduction (and harmonized Highlights wording) to improve readability, flow, and precision without changing scientific claims or conclusions.

#### 3. Point-by-Point Responses

##### Reviewer A

1. **Comment:** *The manuscript by Gildenblat et al describe an innovative approach to interrogate high content spatial mass spectrometry data. This is a timely and well conducted study describing innovative methods and show exciting data illustrating the potential of MALDI MSI. Only minor points remain that should be addressed:*

   **Response:** We thank the reviewer for the very encouraging feedback and have addressed the comments below.

2. **Comment:** *Please specify all abbreviations in figure legends (e.g., SUM, APN, ZI in Fig. 14).*

   **Response:** Thank you. We have revised all figure legends to define abbreviations at first use, including SUM, APN, and ZI (former Fig. 14, now Fig. 12), and ensured consistency across all figure captions.
   *Location in revised manuscript:* Figure legends throughout Results and Supplementary figures.

3. **Comment:** *The authors used a TIMS instrument. Was the ion mobility information used for feature detection?*

**Response:** Thank you for raising this important point. We now explicitly state in the Methods that ion mobility information was calibrated during acquisition but not used for feature extraction or downstream analysis.
*Location in revised manuscript:* Discussion (limitations paragraph addressing TIMS-TOF context) and Methods clarification.

4. **Comment:** *Please cite related MSI in AD studies: PMID: 37006756, 36856542, 34882796, 38299453, 40274785, 40890115.*

   **Response:** Thank you for these helpful references. We have incorporated the suggested literature into the revised manuscript and integrated the citations into the Introduction and Discussion where they provide relevant context for AD-related MSI findings.
   *Location in revised manuscript:* Introduction and Discussion (AD context and ganglioside/plaque interpretation).

#### Reviewer B

##### General Comments

1. **Comment:** *This manuscript presents a computational atlas of mouse brain lipids generated solely from mass spectrometry imaging (MSI) data, leveraging machine learning techniques rather than traditional alternative modalities. The authors present the Computational Brain Lipid Atlas (CBLA) which is a visual framework that enables detailed annotation, dimensionality reduction, and exploration of lipid distributions across hundreds of regions in a mouse brain. Comprehensive molecular visualizations are produced to discover region-specific lipid signatures and disease networks. The workflow is demonstrated by mapping the molecular composition and spatial arrangement of Aβ plaques, revealing their origins and interactions with surrounding tissue. The authors do a great job providing a unique and sound computational and machine learning workflow to discover new connections in a complicated biological system (brain) with highly complex and multiplexed mass spec datasets (MSI). These methods will ultimately advance the interpretation of MSI data and largely contribute to the MSI field. While just focusing on brain lipids here, this framework can be applied to other omics datasets and other organ systems to continue to increase the impact of this work in the future. Additionally, ion images are represented here as a visual DAB IHC stain, which is a neat strategy that I have never seen before, and would be really impactful for those teams that collaborate with pathologists on a routine basis. While the manuscript is thorough, it is recommended that the authors expand on the discussion of the limitations of the workflow, tentatively identify lipids to improve on explaining the actionable biological insight of the model, streamline the main text/figures, and improve on some formatting errors. Following the address of these minor revisions, I recommend publication of the manuscript to Free Neuropathology.*

   **Response:** We thank the reviewer for the encouraging feedback and constructive suggestions. We expanded the Discussion of limitations, added tentative lipid identifications, streamlined main-text figures, and corrected formatting issues.
   *Location in revised manuscript:* Discussion; Supplementary Table S3; revised main and supplementary figure set.

2. **Comment:** *I know the authors have suggested this workflow can be applied to other omics classes, but have you looked at using different matrices to ionize other lipids that could change the visualizations in the model? Why did you stick with NEDC*

   **Response:** Thank you for this important question. We now state why NEDC was selected and note that alternative matrices may capture complementary molecular classes; we also include this as a future direction.
   *Location in revised manuscript:* Discussion (workflow-level limitations and future directions).

3. **Comment:** *In the limitations portion of the discussion, the authors need to discuss more if the model has biases. Does the workflow take into account isotopes and adducts that could be essentially duplicating and weighting specific lipids? How do other ions that are not lipids (chemical interferents on/off tissue or non-lipid endogenous molecules) contribute to the output of the model?*

   **Response:** Thank you for this important point. We expanded the limitations section to address isotopic/adduct redundancy and non-lipid/interferent ions (including off-tissue signals), and we clarify how these factors can affect model weights and outputs and how future filtering may mitigate them.
   *Location in revised manuscript:* Discussion (workflow-level limitations paragraph).

4. **Comment:** *Was spraying an internal standard on the section considered for a more accurate normalization strategy rather than relying on TIC?*

   **Response:** Thank you for this important question. We now explicitly discuss normalization choices, including our rationale for TIC normalization and how internal-standard-based normalization could improve quantitative robustness in future studies.
   *Location in revised manuscript:* Methods (TIC usage statements) and Discussion (normalization limitations/future work).

5. **Comment:** *Are there plans to incorporate additional imaging modalities to bolster the visualizations even more? If so, I would include that in the discussion as a future direction. There are other algorithms, like image fusion (https://pubmed.ncbi.nlm.nih.gov/25707028/), that could still leverage microscopy modalities and build upon your powerful work here.*

   **Response:** Thank you for this helpful suggestion. We have added a paragraph in the Discussion outlining multimodal integration and image-fusion approaches as a natural extension of the current framework, including the referenced strategy (PMID: 25707028).
   *Location in revised manuscript:* Discussion (workflow-level limitations and future multimodal integration).

6. **Comment:** *What is the reason why some of the highlighted m/z's throughout were not attempted to be identified, at the very least by accurate mass let alone MS/MS? You claim these are lipid signals (I believe you based on the mass defect), but don't offer any evidence that this is indeed the case. One suggestion would be to include preliminary/tentative IDs in a supplementary table for the m/z's with ion images shown. I would suggest expanding on the fact that m/z 1179 is detected in the plaques. Numerous papers have shown this and identified the lipid as a ganglioside (GM3 36:1) in mouse models of Alzheimer's disease. This evidence and corresponding citations*

*would add validity to your workflow. Does the actionable insight from your model correspond to what is known about ganglioside biology? The answer to this could drastically increase the impact of the manuscript*

**Response:** Thank you for this important comment. We agree that tentative identifications improve interpretability. We now provide putative annotations for highlighted m/z values (where feasible), expand the discussion of m/z 1179.7308 in relation to reported ganglioside species (e.g., GM3 36:1), and clarify how our spatial patterns are consistent with known plaque ganglioside biology.
*Location in revised manuscript:* Discussion (biochemical interpretation paragraph) and Supplementary Table S3 (putative annotations).

7. **Comment:** *Although manuscript is highly thorough, and journal does not seem to have a figure limit, readability can be improved by moving some non-essential figures to the supplementary. It is my belief that just because you CAN include 16 figures, doesn't always mean you SHOULD. The authors should prioritize what the reader should take away from the manuscript and I believe having 16 main figures seriously dilutes that priority.*

   **Response:** Thank you for this helpful suggestion. We have revised the figure organization by moving selected non-essential figures and display panels to the Supplementary Material, improving narrative focus and readability while preserving full data accessibility.
   *Location in revised manuscript:* Main text figure flow and Supplementary Figures S1–S2.

##### Detailed Comments

1. **Comment:** *Title: I don't know if you need the "-" between mass spectrometry and I would suggest adding "mouse brain" to title. Other than that, good and descriptive title.*

   **Response:** Thank you for this helpful suggestion. We have revised the title wording to include explicit mention of the mouse brain context and adjusted the formatting accordingly.
   *Location in revised manuscript:* Title.

2. **Comment:** *Abstract: "A new algorithm decomposes annotated regions into precise m/z features and resolves full-precision m/z values from MSI data." Need to explain this better or use other verbiage in the abstract. I did not know what a "full-precision m/z value" was until much further into the manuscript. From my understanding it's an alignment processing method so I would recommend clarification early on.*

   **Response:** Thank you for this helpful suggestion. We have rewritten this sentence in the abstract to clarify the concept and avoid ambiguity, replacing the term "full-precision m/z values" with clearer terminology ("high-resolution m/z values") and ensuring consistency throughout the manuscript.
   *Location in revised manuscript:* Abstract.

3. **Comment:** *Introduction: "The actual problem here lies in the poor presentation of the MSI data as ion images and the actual lack of morphological expertise." I believe this is the comparison statement that justifies your new workflow/technology, and needs to be explained more and expanded on to really emphasize the purpose of your mansucript. The following sentence after*

*this one seems to be written in a manner that highlights the advantages of your workflow, but this isn't clear based on the sentence structure. You then go into explaining other challenges associated with different workflows to highlight the advantages of yours once again. I recommend fixing the organization of the thought process across these two paragraphs because it clouded my interpretation of your message.*

**Response:** Thank you for this important comment. We have restructured the relevant paragraphs in the Introduction to improve logical flow and clarify the motivation for the proposed workflow. In particular, we revised the problem statement and its supporting arguments to more clearly distinguish between limitations of existing approaches and the advantages of our method. We also performed sentence-level revisions across the Introduction to improve clarity, consistency, and phrasing while preserving the scientific meaning.
*Location in revised manuscript:* Introduction (opening rationale paragraphs).

4. **Comment:** *Introduction: You should remove exclamation point after "modalities"*
   **Response:** Thank you, we have removed the exclamation point.
   *Location in revised manuscript:* Introduction.

5. **Comment:** *"We demonstrate the power of this new pipeline by analyzing Aβ plaques and, for the first time, quantify their molecular relationship with surrounding brain regions, demonstrating that Aβ plaques incorporate specific lipid signatures originating from affected cerebral cortex regions." I would strongly suggest to stay away from the idea/word of "quantifying" because it begins a debate of relative vs absolute in mass spectrometry data. I know you suggested a future strategy would be to use quantitative scoring for the annotations, which is a good idea, but be cautious specifically of the quantification terminology.*

   **Response:** Thank you for this important suggestion. We agree that the term "quantifying" may be misleading in the context of MSI data. We have revised the sentence to avoid this terminology and instead use "characterizing" to more accurately reflect the nature of the analysis, while also improving clarity and precision of the statement.
   *Location in revised manuscript:* Introduction and Abstract wording updates.

6. **Comment:** *Methods: "We used the machine standard of 20 × 20µm2 (4 × 4 pixels of 5 × 5µm2 ) to generate sufficient signal (AU) also for very low abundant species. 5 × 5µm2 generates higher spatial resolution for the cost of the loss of low signals." Fix grammar mistakes throughout.*

   **Response:** Thank you for this helpful comment. We have corrected grammar and style throughout the Methods, including revising the cited sentence on acquisition settings and signal trade-offs to improve clarity and readability.
   *Location in revised manuscript:* Methods (acquisition settings paragraph) and global language edit.

7. **Comment:** *Figure 1- Better caption is needed or put labels on the visualizations. Are they different sections of different brains? Can you fit any large annotations in the figure?*

   **Response:** Thank you for this helpful suggestion. We have expanded the Figure 1 legend, added clarifying labels to indicate section and sample context, and improved annotations within the

figure to enhance interpretability.
*Location in revised manuscript:* Figure 1 legend and panel labeling.

8. **Comment:** *"m/zs" is used throughout. Make sure all of them are "m/z's" and italicize m/z*

   **Response:** Thank you for this helpful comment. We have standardized notation throughout the manuscript, ensuring consistent *m/z* formatting and italicization (*m/z*) across the text and figures.
   *Location in revised manuscript:* Main text, figure legends, and Supplementary Material.

9. **Comment:** *Results: "This feedback loop helped us improve the quality and granularity of our initial annotations, as well as detect previously unknown subnuclei, such as those in the Substantia nigra and the Globus pallidus." Revealing a biological unknown is a very significant claim. Is there any biological pretense or corroborating evidence for this? If not, I would soften the language by using words like "suggesting"*

   **Response:** Thank you for this important comment. We have revised this statement to soften the language and now use wording that reflects suggestive rather than definitive evidence, as independent biological validation is not yet available.
   *Location in revised manuscript:* Results and Discussion phrasing of biological interpretation.

10. **Comment:** *Figure 8A- This is the first ion image that made me question what is on tissue vs off tissue. Yes the m/z 480 could be on the outer most region of the section, but it appears to be heavily abundant off-tissue. Is there an explanation for this and how it influences the output of the model (similar to general comment above)? You may want to include something in the supplement where your ion images are overlaid onto the brain annotations; similar to Figure 11, but rather side by side with an H&E, you would include a direct registered semi-transparent overlay of the brain map and ion image. An alternative solution would be to add a dotted line to ion images to delineate on and off tissue.*

    **Response:** Thank you for this important comment. We added text on the origin and interpretation of off-tissue signals, clarified potential effects on model outputs, expanded the limitations discussion, and improved supplementary overlays/annotations to delineate on- versus off-tissue regions.
    *Location in revised manuscript:* Discussion (workflow-level limitations) and revised supplementary visualization panels.

11. **Comment:** *Figure 12- Are the ion images (iii-vi) for serial sections throughout a single brain? Some other figures (13,14) are like this as well (Figure 14 explains it better). Please state these details in each figure caption because it is not clear to me, unless I am missing something.*

    **Response:** Thank you for this helpful comment. We have revised the figure captions to explicitly clarify section provenance, including whether images correspond to serial sections from a single brain, and ensured this information is consistently stated across all relevant figures.
    *Location in revised manuscript:* Figure captions for multi-section image panels (including former Figs. 12–14 context).

12. **Comment:** *Figures 10 and 13- The CBLA is so small to read in these figures. I would suggest improving formatting in some manner or move to the supplement to increase the size.*

**Response:** Thank you for this helpful suggestion. We improved readability by moving non-essential explanatory panels to the Supplementary Material and enlarging the main figures. Former Figure 10 is now Supplementary Figure S1, and former Figure 13 is now Supplementary Figure S2; the corresponding main figures (now Figures 8 and 11) were reformatted and enlarged.
*Location in revised manuscript:* Main text figure revisions and Supplementary Figures S1–S2.

13. **Comment:** *Discussion: Expand this section based on the general comments above.*

    **Response:** Thank you for this suggestion. We have expanded the Discussion to address the requested methodological limitations, validation considerations, and future directions.
    *Location in revised manuscript:* Discussion (limitations and future directions paragraphs).

14. **Comment:** *Discussion: Last paragraph of discussion is a duplicate of the conclusion so you need to delete one.*

    **Response:** Thank you for noting this. We have removed the duplicated text at the end of the Discussion.
    *Location in revised manuscript:* Discussion ending.
